# *Plasmodium* Repetome: A mysterious space with a wealth of information

**DOI:** 10.1101/2025.04.14.647515

**Authors:** Satinder Kaur, Vikash Kumar, Karanbir Singh, Ankita Behl, Arpna, Prakash Chandra Mishra, Rachna Hora

**Author notes:** Correspondence: Rachna Hora; E mail; Contact No. Extn: 3399; Prakash Chandra Mishra, Contact No.: Extn: 3220. Equal contribution (Satinder Kaur, Vikash Kumar).

## Abstract

Eukaryotic proteomes harbour repetitive stretches of amino acids that may play critical roles in the biology of that organism. While several tandem repeats (TR) are known to contribute to protein structure and function, information about the vast majority of repeat regions remains obscure. In this article, we have analysed the repeat content of different *Plasmodium* species and found the leading human malaria-causing *P. falciparum* (Pf) and *P. vivax* to be exceptionally rich in TR regions (>40% TR containing proteins). Detailed analysis of Pf ‘repetome’ showed this intracellular parasite to carry longer TRs, several of which were present in exported proteins important for parasite survival and immune evasion. The repeat regions of Pf were enriched in acidic amino acids and asparagine (Asn), where Asn was more abundant in short and intermediate TRs, suggesting an evolutionary bias influenced by replication slippage and positive selection. Gene ontology analysis of TR containing Pf proteins helped us to understand their cellular localization along with the molecular and biological processes they are involved in. The Pf variable surface antigen families with roles in important pathogenic processes like cytoadherence, immune evasion *etc.* had low repeat content present within seroreactive peptides. Three-dimensional structure predictions of TR regions revealed several repeats to adopt ordered super-secondary conformations that are known to facilitate intermolecular interactions. Overall, this is a comprehensive study attempting to gain insights on the importance of TRs in malaria parasite biology and suggests a novel route to understanding protein function through the characterization of repeat content.

## Introduction

Tandem repetitions of peptidic sequences frequently occur in proteomes, where they may contribute to the structure or function of these proteins [1], [2]. While tandem repeats (TR) are present in approximately 14% of all proteins, occurrence of amino acid TRs is three times more likely in eukaryotes [3]. These ubiquitous modules range from single amino acid repetitions to repeats of oligomeric or polymeric units that may span a hundred or more residues [4], [5]. Plausible mechanisms that bring about TR occurrence include replication slippage within DNA microsatellite regions, or DNA recombination and duplication events for the longer minisatellite and satellite regions [6], [7], [8]. Historically believed to be translated from junk DNA and hence non-functional, repeat units in proteins are now increasingly reported to be involved in generation of immune responses, cell signalling, transcriptional regulation and interactions with other proteins, nucleic acids or metal ions *etc.* [9], [10], [11], [12], [13], [14], [15]. Such regions display dynamic contractions and expansions leading to rapid alterations in genomic DNA that may modify activity of the expressed protein or even impart a novel function to it [16], [17]. Amino acid repeats are reported to carry particular significance in pathogenesis of different diseases e.g. function of poly Q stretches in Huntington’s disease, role of parasite surface proteins TRs in invasion *etc.* [18].

Oligomeric amino acid TRs are particularly widespread among several parasitic organisms including *Plasmodium* species [19]. The percentage of TR genes was the highest in *Plasmodium falciparum* (Pf) (3.07%) when compared with other protozoan parasites (<1%) like *Leishmania infantum, Leishmania major, Trypanosoma brucei and Theileria annulata* [20]. Pf is particularly rich in homo-polymeric asparagine/ glutamine tracts that are present in more than 24% of its proteins [21]. Such stretches are associated with prion-like domains responsible for protein aggregation at elevated temperatures. Intracellular *Apicomplexan* parasitic protozoans like *Plasmodia, T. gondii* and *E. tenella* showed higher repetition in their proteins when compared with extracellular parasites and free living protists [22]. Most repeat-containing proteins are either secreted or localized to the parasite surface while being enriched in residues encompassing glycosylation sites [23]. Repeats contained within intracellular parasite proteins are predominantly perfect, while those of extracellular parasites show more degeneracy [22]. Proteomes of eukaryotic parasites display a significantly better correlation between their percentage of repetitive amino acids and average protein length as compared with free-living eukaryotic proteomes. Repeat regions have therefore been hypothesized to have roles in parasite adaptation in diverse ecological niches and immune evasion through host-parasite interactions during invasion of host cells and parasite life cycle. TRs are also reported to occur in toxins and virulence factors of pathogens [24].

*Plasmodium falciparum* is a parasitic protozoan that expresses over 5000 proteins [25]. Several proteins of this malaria causative organism are reported to carry TRs that perform functions crucial for parasite survival and disease pathogenesis [17]. A prominent example is the ‘NANP repeats’ of circumsporozoite protein (CSP) that constitute the immunodominant B-cell epitopes of the protein [26]. Deletion of the CSP repeat region in parasite lines led to defective development of sporozoites [27]. The malaria vaccine RTS,S is therefore based on this central repeat region of CSP [28], [29]. Yet another example highlighting the significance of repetitive sequences in Pf is the ‘C-terminal lysine rich repeat sequence’ of KAHRP (knob-associated histidine-rich protein), which forms knob like structures on the surface of infected red blood cells (iRBCs) and harbours PfEMP1 (Pf erythrocyte membrane protein 1), the key virulence factor responsible for cytoadherence. Deletion of the repeat region of KAHRP resulted in smaller knob formation and reduced cytoadhesion of iRBCs [30]. Therefore, repeat regions of different Pf proteins are known to be involved in crucial pathogenic processes and manifestations like cytoadherence, heme detoxification, genesis of Maurer’s clefts, localization of proteins and cellular rigidity [17], [31], [32], [33], [34], [35], [36]. Binding of repeats with proteins, nucleic acids, metal ions, lipids *etc.* makes significant contribution to the above functions [37]. Patient antibodies with reactivity against repeat regions of CSP, SERA5 (serine rich antigen 5), MSP1 (merozoite surface protein 1) and GLURP (glutamate rich protein) had the ability to cause parasite growth inhibition, emphasizing the importance of TRs in generation of the host immune response [38], [39], [40], [41]. It has however also been hypothesized that repetitive amino acid sequences create an ‘antigenic smokescreen’ that obstruct affinity maturation of protective antibodies [19].

Low complexity regions (LCRs) are usually unstructured domains but some can take up a structured organization [3], [42], [43]. It is commonly understood that proteins that show structural convergence despite dissimilarities in their primary sequences are likely to perform common functions [44]. RepeatsDB is a database that contains a set of predicted three-dimensional structures of structured tandem repeat proteins (STRPs) [45]. STRPs are classified on the basis of the length of their repeat unit and arrangement *viz.* Class I: ‘crystalline aggregates’ of mono or di amino acid repeats, Class II: ‘fibrous structures’ with repeat units of 3-4 residues (collagens and alpha helical coiled coils), Class III: ‘elongated structures’ with TR length ranging from 5 to 40 amino acids (solenoid and non-solenoid structures), Class IV: ‘closed structures’ having ∼20 to 60 amino acid long TRs (TIM barrel, closed solenoid, β propeller *etc.*) and Class V: ‘beads on a string’ that have independently folding repeat units of >50 amino acids [2], [46], [47]. Repeats of intermediate length (20–50 amino acids) are most commonly observed, which may serve to form integrated assemblies that provide multiple binding sites (e.g. collagens, myosin, keratins etc) [1]. This feature may serve to recruit multiple copies of an interacting partner to enhance avidity of the interaction [17]. Also, highly connected proteins involved in formation of interaction hubs in cells are often rich in repetitive content but lack structure [48]. Such intrinsically disordered repeat regions may however tend to assume an ordered conformation when bound to a partner [49]. Interestingly, TR regions tend to be less structured when comprised of perfect repeats [50].

In this article, we have systematically analysed the proteome sequences of ten different *Plasmodium* species available on PlasmoDB to evaluate the abundance of their repeat content termed ‘repetome’. The distribution of repetitive sequences in different *Plasmodia* has also been studied as a function of the repeat unit length (period). We have investigated the repeats of the most fatal *Plasmodium* species i.e. *P. falciparum* in further detail to understand the amino acid composition and bias of the Pf repetome. Gene ontology analysis was used to understand whether repeats are associated with proteins having specific roles. Our article also evaluates the repeat content of Pf variable surface antigen (VSA) families, which are critically involved in disease pathogenesis and elicitation of host immune responses. We have further attempted to analyse the predicted structures of a few prominent TR domains from Pf proteins for their secondary structure content and potential to form super-secondary structures. Super-secondary structures adopted by TR regions of a handful of such proteins were used to predict their probable functions. This article provides a comprehensive overview of the repetome of different *Plasmodium* species, particularly *P. falciparum*, and offers fresh insights into the structure-function relationships of this black box of parasite biology.

## Materials And Methods

### Data Retrieval

The protein sequences of ten different *Plasmodium* species (including the ones encoded by the mitochondrial and apicoplast genes) were downloaded from PlasmoDB and saved as .txt files [51] [52]. Sequences were retrieved for different *Plasmodia* causing human, simian and rodent malaria. The species and strains studied (number of proteins in respective proteome given in brackets) were *P. falciparum* 3D7 (5318), *P. vivax* Sal-1 (5530), *P. malariae* UG01 (5942) & *P. knowlesi strain H* (5328) known to infect humans, *P. gaboni G01* (5134), *P. reichenowi G01* (5644) & *P. cynomolgi strain B* (5716) that infect simians and *P. berghei ANKA* (4945), *P. yoelii yoelii 17XNL* (7724) & *P. chabaudi chabaudi* (5199) that cause rodent malaria. Protein sequences corresponding to the longest transcript were retrieved for genes where multiple transcripts were available.

### Prediction of tandem repeats

Tandem repeats in the downloaded protein sequences were identified by using XSTREAM software, which is a JAVA encoded tool for TR identification in FASTA sequences (Supplementary file 1) [53]. The command line version of XSTREAM was downloaded from https://amnewmanlab.stanford.edu/xstream/in on a local computer. XSTREAM was run by executing the command ‘java -Xmx1000m -Xms1000m -jar xstream.jar myFile.txt’ for its default settings. The default parameters of XSTREAM are set to identify TRs in protein sequences with minimum word match 0.7 (>70% identity amongst repeat units), minimum consensus match 0.8 (>80% identity with the consensus sequence), minimum copy number 2 (number of repetitions of repeat unit), minimum period 3 (length of repeat unit), minimum TR domain length 10 (total length of the identified repeat region), maximum consecutive gaps 3 (number of amino acids inserted between successive repeat units) and maximum indel error 0.5 (number of gaps in TR domain/ TR domain length). The arguments -m2 and -z were included in the command to change the minimum period to 2 and create an excel spreadsheet of the TR output respectively. The above parameters for identification of repeats provide moderate TR degeneracy with a significant P value of <0.02. The obtained data were plotted and analysed as follows. 1. The proportion of TR containing proteins in the proteomes of different *Plasmodium* species. 2. TR usage in different *Plasmodium* species, which is calculated as a ratio of the ‘total number of proteins in a proteome’ and the ‘number of TR containing proteins in that proteome’. 3. Percentage of TR containing proteins in the proteome of *Plasmodium falciparum* plotted as a function of period lengths 2, 3, 4, 5, 6-10, 11-15, 16-20, 21-30, 31-100 and 101-200. The number of TR proteins in the Pf proteome was also tabulated 4. Usage of each amino acid (number of times used and percentage) in total identified tandem repeats (276023 amino acids) of Pf. 5. Occurrence of each amino acid in repeat regions of Pf relative to the complete proteome (4064438 amino acids). Relative occurrence is determined by calculating the difference in the ‘usage of each amino acid (expressed as percentage) in the repeat regions’ of Pf and the ‘amino acid usage in the entire Pf proteome’. ProtParam tool on web.expasy.org was used to count the number of times each amino acid was present in the TR region and whole proteome of Pf [54]. 6. Considering the asparagine (Asn) richness in Pf proteome, Asn usage trend in TRs was determined and analysed as a function of the period length. The selected periods were 2-5, 6-10, 11-15, 16-20, 21-30, 31-100, 101-200 and 200-300. The degree of Asn usage bias was calculated by using the following formula-

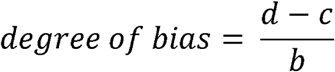

b = Total number of amino acids in TR region of given period length

c = Expected number of Asns in each period group in the absence of usage bias

d = Number of Asns used in each period group of TR regions

### Functional characterization of repeat containing Pf proteins

The TR containing proteins of Pf were subjected to gene ontology (GO) analysis. For these analyses, TR identification using XSTREAM was repeated while fixing the search parameters to copy number 2, period 3 and minimum TR domain length 20. A comprehensive datasheet for the above identified proteins was retrieved from PlasmoDB (plasmodb.org). Lists of PlasmoDB IDs of the TR containing proteins and their corresponding GO terms for different classes (cellular component, molecular function and biological processes) were run through TB tools to obtain the number of proteins placed under each GO term [55]. These proteins were categorized manually on the basis of their GO classes to broadly understand the functions of repeat containing Pf proteins. The results were represented in a pictorial form. A subset of TR containing proteins associated with Enzyme Commission (EC) numbers were listed separately and analysed by placing them into the seven different enzyme classes (oxidoreductases, transferases, hydrolases, lyases, isomerases, ligases, translocases).

Another subset of proteins specific to *Plasmodium* species comprising the Variable Surface Antigen (VSA) protein families (SURFINS, PfEMP1, RIFIN, PfMC-2TM, STEVOR) were also listed separately for assessing their TR content by XSTREAM analysis. Total number of members in each family were determined from this list followed by the omission of pseudogenes. The parameters used for the XSTREAM run were set to copy number 2, period 3 and minimum TR domain length of 20. Results of the XSTREAM run were analysed to determine the number and percentage of proteins containing TRs along with the lowest and highest copy number and period. The identified repeat regions were compared with the seroreactive proteins enlisted by Raghavan *et. al.* to understand the potential of repeat motifs to act as epitopes [56].

### Structural characterization of TR containing Pf proteins

TR regions from Pf proteins spanning all periods and having highest copy number were selected to assess their structures. The protein data bank (https://www.rcsb.org/) and AlphaFold protein structure database (https://alphafold.ebi.ac.uk/) were searched for available structures of the selected proteins [57], [58], [59]. For AlphaFold structures of TRs having very low (pLDDT <50) to low confidence (pLDDT <70), the three-dimensional structures were predicted by using the SWISS-MODEL (https://swissmodel.expasy.org/) [60]. TR regions predicted to adopt an ordered conformation were analysed further for their structural and functional characteristics.

### Structure - function analysis of TR containing proteins

TR regions with ordered structures were assessed for their percentage coverage in the respective protein, periodicity, copy number and type of secondary/ super-secondary structure adopted. TR regions forming super-secondary structures were further analysed to identify structurally similar domains by using NCBI-VAST (vector alignment search tool) (https://www.ncbi.nlm.nih.gov/Structure/VAST/vastsearch.html) [61], [62]. The best VAST hits were then used to identify the structural fold taken up by the TR being studied by using PDB database. Four proteins having super-secondary folds with repetitive patterns were selected to understand their structure and function better. Here, the first three NCBI-VAST hits having the best VAST scores were used for analysis.

## RESULTS

### *P. vivax* and *P. falciparum* have the highest tandem repeat share

Tandem repeats (minimum period 2, minimum copy number 2, minimum TR domain length 10) were identified in ten different *Plasmodium* species by using XSTREAM software. XSTREAM allows large scale sequence analysis for identification of perfect and degenerate repeats. Most *Plasmodium* species included in our study contained more than 3000 tandem repeat units, ranging from 1504 in *P. berghei* to 6928 in *P.vivax* (Table 1). *P. berghei* and *P. vivax* also contained the least (857) and highest (3731) number of unique TRs respectively.

**Table 1:**
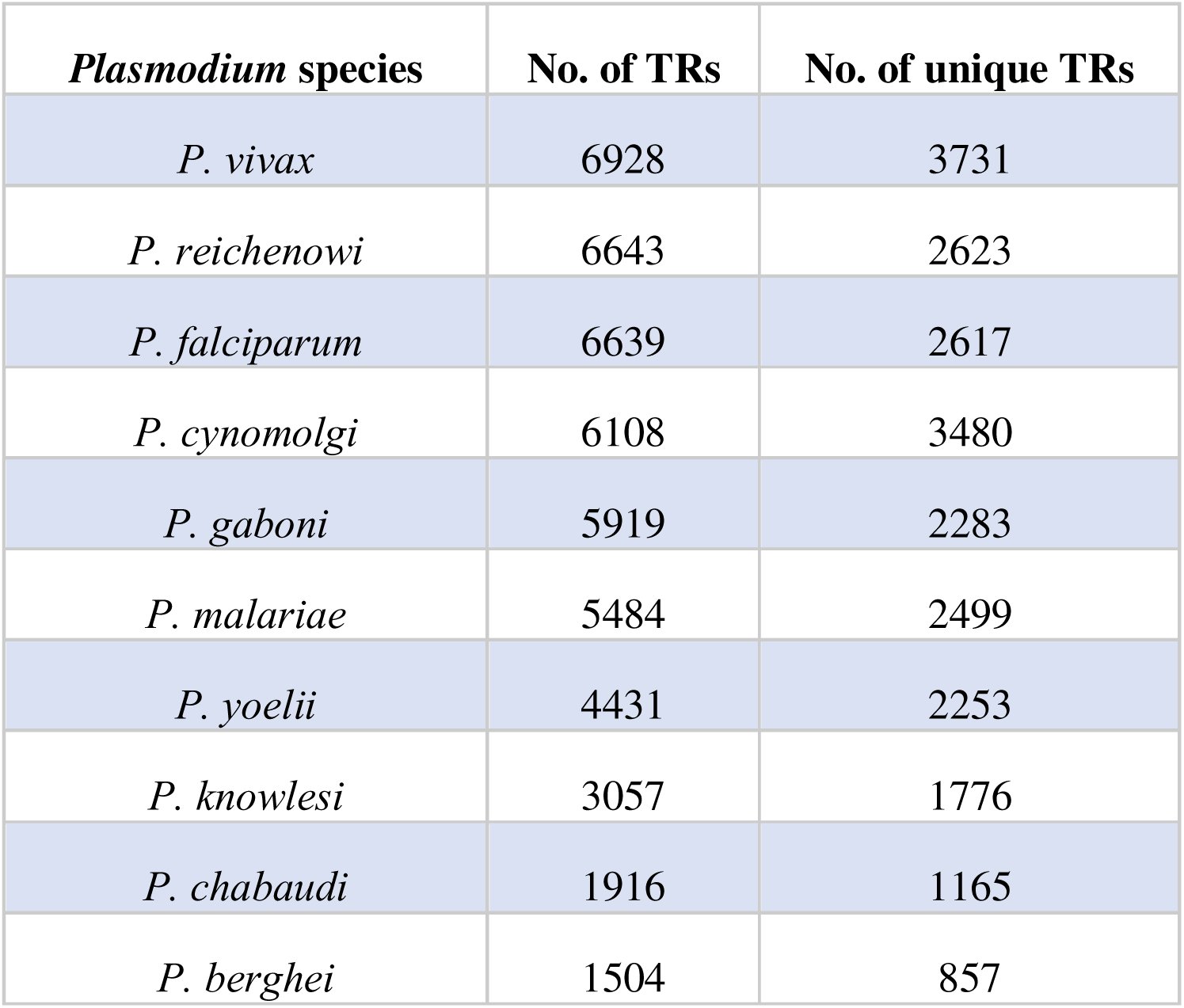
Number of oligomeric TR units and unique TRs in Plasmodium species: Table shows the total number of tandem repeats and unique TRs in ten *Plasmodium* species as identified by XSTREAM (Copy number 2, period 2 and minimum TR domain length of 10).

| ***Plasmodium* species** | **No. of TRs** | **No. of unique TRs** |
| --- | --- | --- |
| *P. vivax* | 6928 | 3731 |
| *P. reichenowi* | 6643 | 2623 |
| *P. falciparum* | 6639 | 2617 |
| *P. cynomolgi* | 6108 | 3480 |
| *P. gaboni* | 5919 | 2283 |
| *P. malariae* | 5484 | 2499 |
| *P. yoelii* | 4431 | 2253 |
| *P. knowlesi* | 3057 | 1776 |
| *P. chabaudi* | 1916 | 1165 |
| *P. berghei* | 1504 | 857 |

The number of proteins carrying TRs in each of the *Plasmodium* species were determined (Figure 1a). The proteomes of human infecting *P. vivax* (Pv) and *P. falciparum* (Pf) were found to carry the maximum proportion i.e. 41.52% and 40.97% of TR containing proteins respectively. These two *Plasmodium* species are responsible for most malaria cases globally. The simian malaria causing species *P. gaboni* (Pg) and *P. reichenowi* (Pr) closely followed with 40.44% and 40.01% TR proteins, while 35.93% proteins of *P. cynomolgi* (Pc) contained TRs. Human infecting species *P. malariae* (Pm) (33.46%) and *P. knowlesi* (Pk) (29.94%) carried a significantly lower proportion of TR containing proteins than their counterparts Pv and Pf. While Pv and Pk are closely related phylogenetically, they carry significantly different proportions of repeats [63]. The least proportion of TR proteins were present in rodent malaria species *P. yoelli* (Py) (27.21%), *P. chabaudi* (Pch) (22.02%) and *P. berghei* (Pb) (20.22%). The percentage of TR proteins was found to be considerably different in the evolutionarily close Py and Pb as well [64].

**Figure 1:**
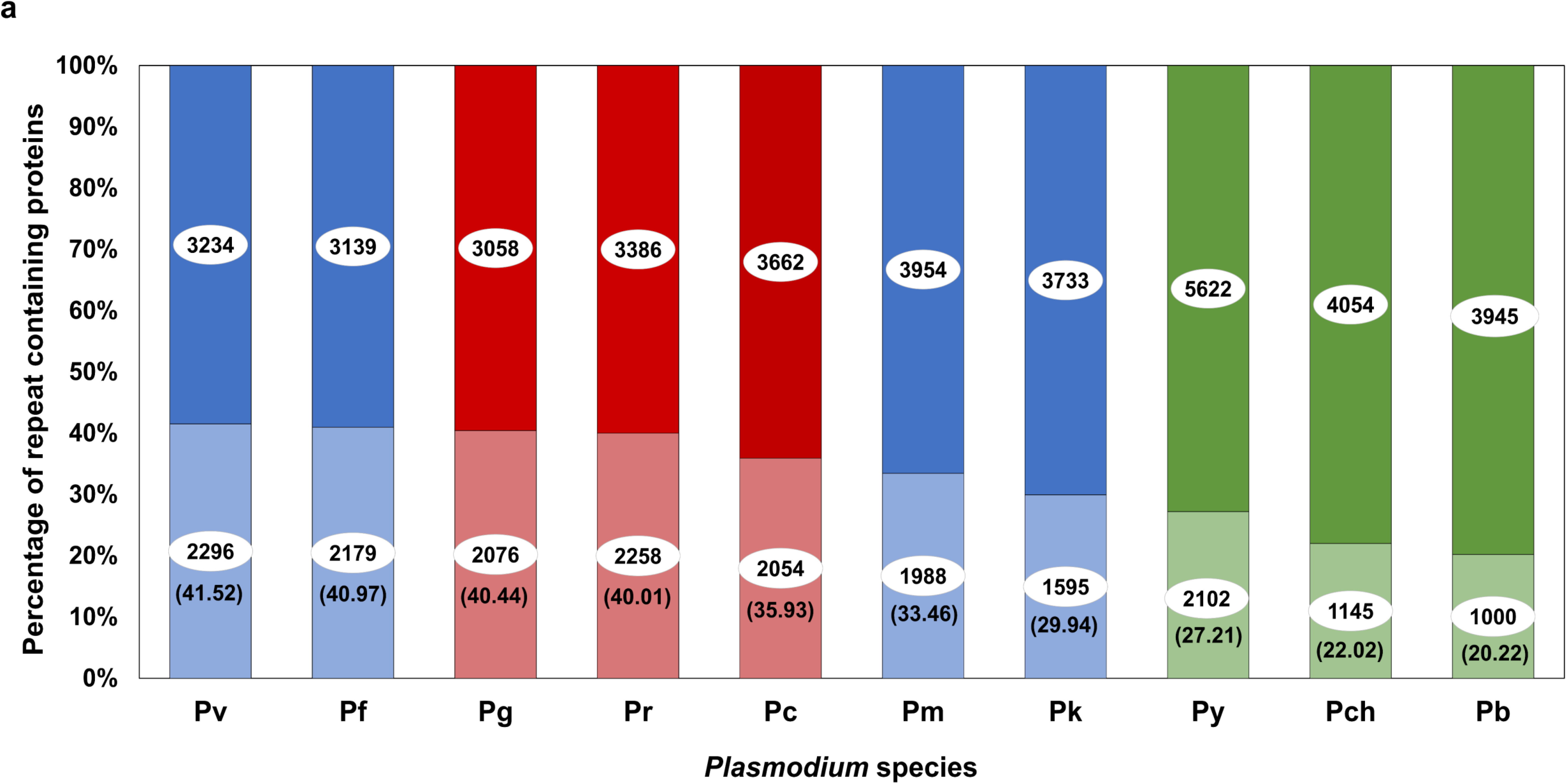
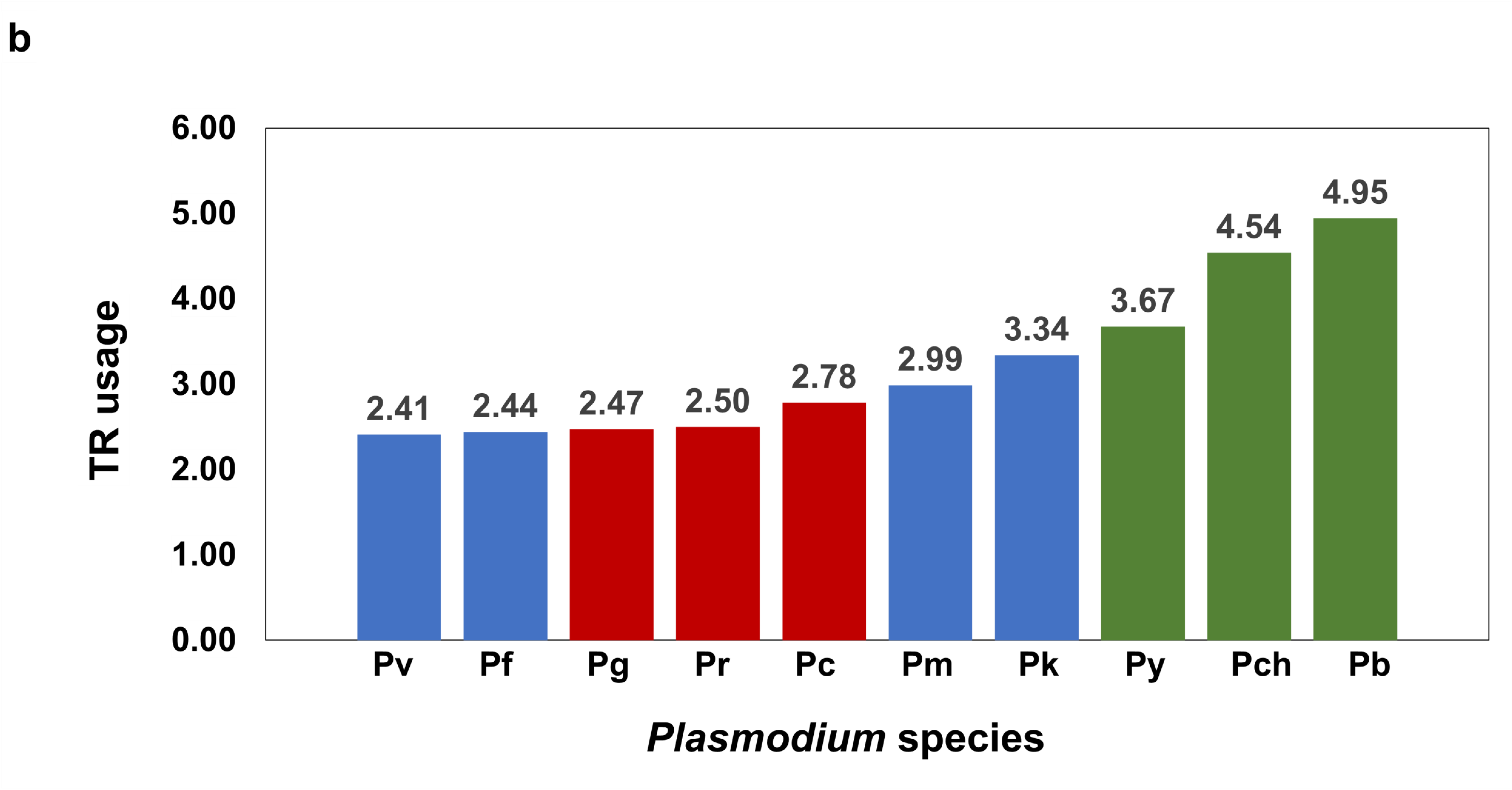
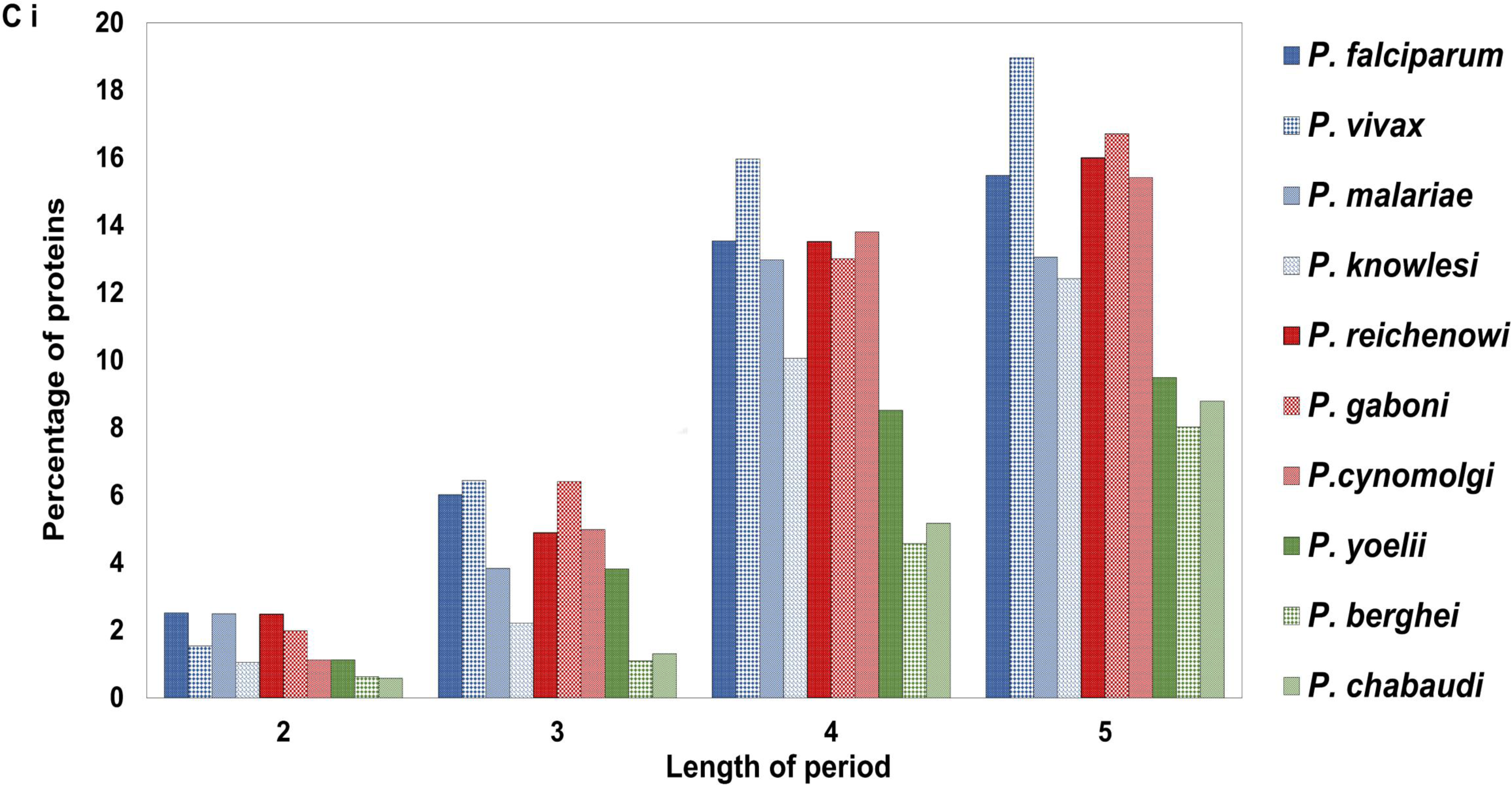
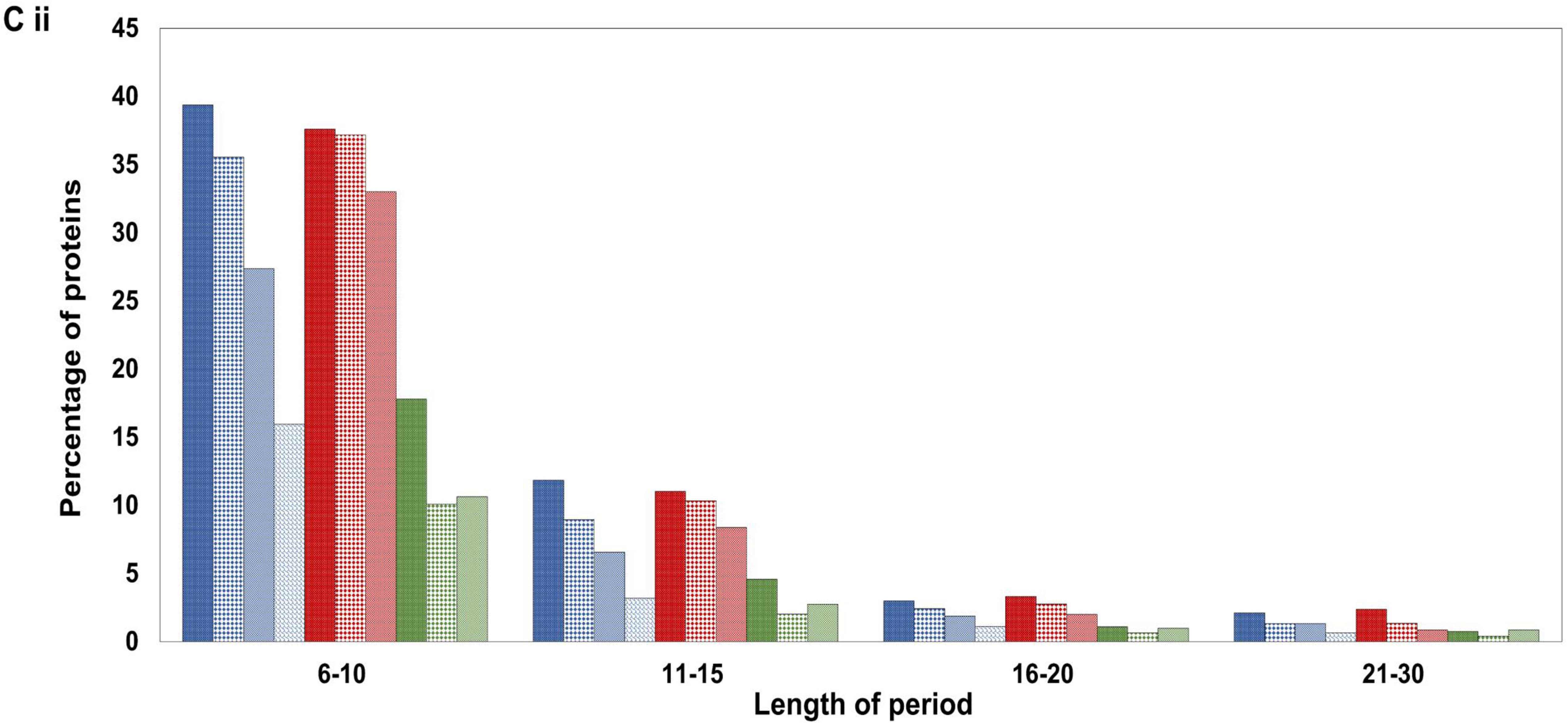
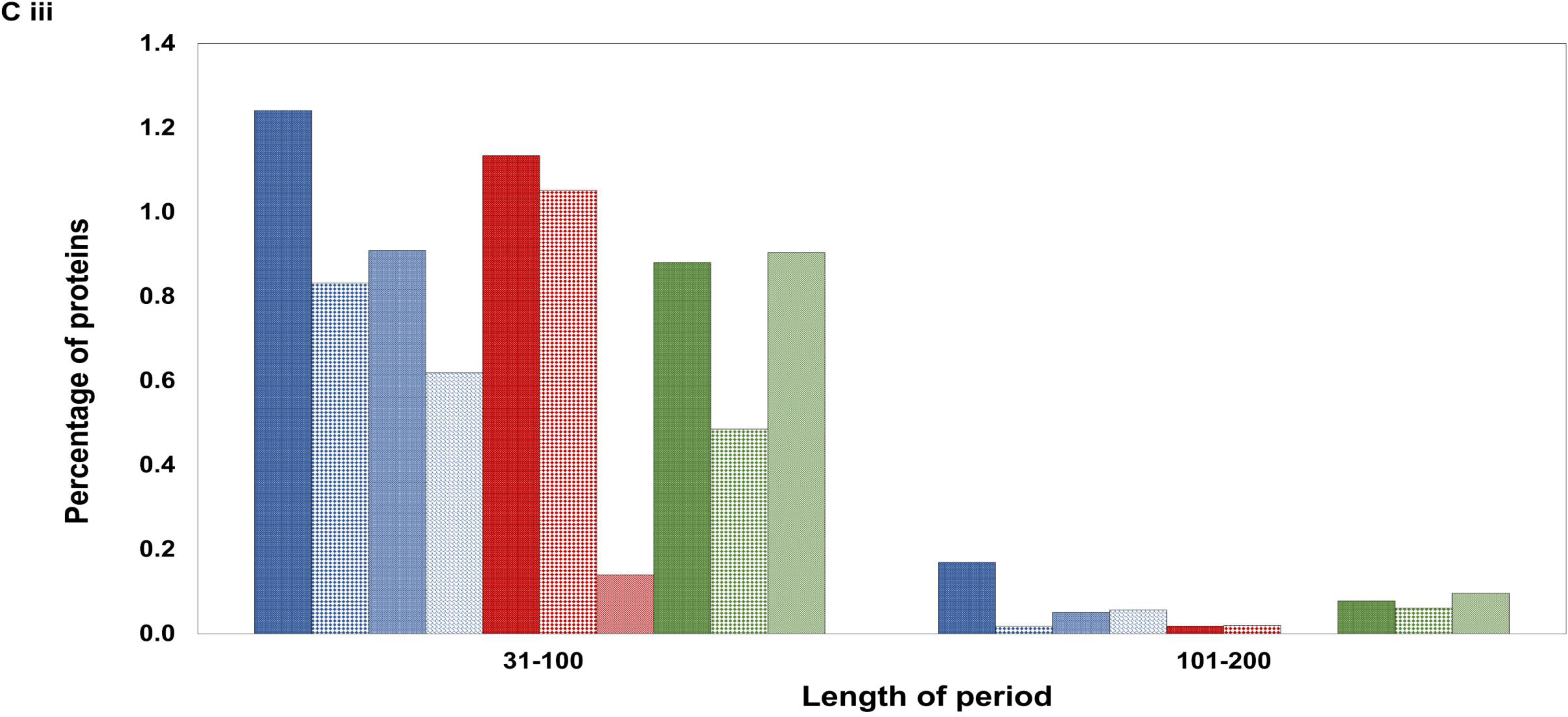
Abundance of repeat-containing proteins in ten Plasmodium species. Bar diagrams showing data for species causing human malaria (blue bars); simian malaria (red bars) & rodent malaria (green bars). Species and strains studied were *Plasmodium falciparum* 3D7 (Pf), *P. vivax* Sal-1 (Pv), *P. malariae* UG01 (Pm), *P. knowlesi strain H* (Pk), *P. gaboni G01* (Pg), *P. reichenowi G01* (Pr), *P. cynomolgi strain B* (Pc), P*. berghei ANKA* (Pb), *P. yoelii yoelii 17XNL* (Py) and P. *chabaudi chabaudi* (Pch). a) Stacked column chart showing the percentage of TR-containing proteins in ten different *Plasmodium* species *vs* the total number of proteins in the proteome considered as 100%. Light shades represent TR-containing proteins and darker shades denote the proteins without repeats. The number of proteins with and without TRs are inscribed in white ovals in the respective area within the shaded columns. The percentage of TR-containing proteins for each species is given in brackets. b) Bar diagram representing TR usage for ten *Plasmodium* species. c) Bar diagram representing the percentage of TR containing proteins (out of total number of proteins in that species) as a function of period (repeat length). c (i) shows percentage of TRs in 2, 3, 4 and 5 mers. c (ii) shows TR percentage in periods 6-10, 11-15, 16-20 and 21-30. c (iii) represents percentage of TRs for repeat lengths 31-100 and 101-200. Species name for Figure 1c is indicated in a panel on figure c (i).

The above data were used to understand TR usage in various *Plasmodium* species (Figure 1b). TR usage was high in Pv, Pf, Pg and Pr where TRs were present in nearly one out of every ∼2.5 proteins. Rodent malaria species Pb had the lowest TR usage with one out of nearly five proteins containing repeats.

### *P. falciparum* proteome is rich in long TRs

We compared the percentage of TR containing proteins in different *Plasmodium* species in a period-wise fashion (Figure 1c and Table 2). Our results found that the period of TRs in various *Plasmodia* ranges from 2 to 411 amino acids. Pv contained the highest percentage i.e. 42.9% of short repeats (2-5 mers) followed by Pg (38%), Pf (37%) and Pr (36.8%). However, Pf carried a greater percentage (39.3%) of 6-10 mers than Pv (35.5%). All *Plasmodium* species except Pf and Pr contained a higher proportion of 2-5 mer repeat containing proteins than 6-10 mer. The evolutionarily close Pf and Pr followed similar patterns with respect to their period-wise TR containing proteins [65]. Both Pf and Pr contained the highest percentage of >10 amino acid long repeats carrying proteins when compared with other species viz ∼11% 11-15 mers, ∼3% 16-20 mers, ∼2% 21-30 mers and ∼1% 31-100 mers. Pf carried the largest number of very long TRs (nine 100-200 mers and two 200-300 mers). Other *Plasmodium* species with relatively higher numbers of very long TRs (>100 amino acid long) include Py having 7 TRs, Pch with 6 and Pm with 5. Both Pm and Pch carried one extremely long repeat of 411 amino acids. Pc was found to have the least number of long repeat units with a maximum period of 51 amino acids.

**Table 2:**
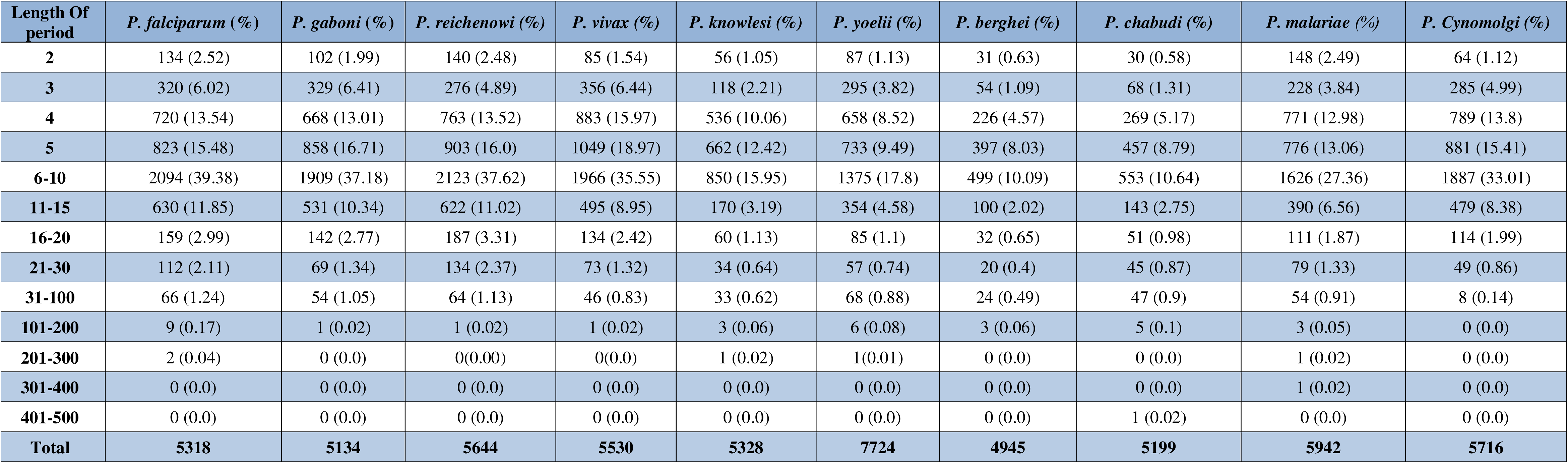
Number of TR-containing proteins as a function of period. Table shows the number of proteins containing TRs of specific periods in different *Plasmodium* species. The considered periods are 2, 3, 4 & 5 and period ranges 6-10, 11-15, 16-20, 21-30, 31-100, 101-200, 201-300, 301-400 & 401-500. Total number of proteins in each *Plasmodium* species is also tabulated. The percentage of proteins (total proteome = 100%) having TRs of a particular period is given in brackets.

| Length Of period | *P. falciparum* (%) | *P. gaboni* (%) | *P. reichenowi* (%) | *P. vivax* (%) | *P. knowlesi* (%) | *P. yoelii* (%) | *P. berghei* (%) | *P. chabudi* (%) | *P. malariae* (%) | *P. Cynomolgi* (%) |
| --- | --- | --- | --- | --- | --- | --- | --- | --- | --- | --- |
| 2 | 134 (2.52) | 102 (1.99) | 140 (2.48) | 85 (1.54) | 56 (1.05) | 87 (1.13) | 31 (0.63) | 30 (0.58) | 148 (2.49) | 64 (1.12) |
| 3 | 320 (6.02) | 329 (6.41) | 276 (4.89) | 356 (6.44) | 118 (2.21) | 295 (3.82) | 54 (1.09) | 68 (1.31) | 228 (3.84) | 285 (4.99) |
| 4 | 720 (13.54) | 668 (13.01) | 763 (13.52) | 883 (15.97) | 536 (10.06) | 658 (8.52) | 226 (4.57) | 269 (5.17) | 771 (12.98) | 789 (13.8) |
| 5 | 823 (15.48) | 858 (16.71) | 903 (16.0) | 1049 (18.97) | 662 (12.42) | 733 (9.49) | 397 (8.03) | 457 (8.79) | 776 (13.06) | 881 (15.41) |
| 6-10 | 2094 (39.38) | 1909 (37.18) | 2123 (37.62) | 1966 (35.55) | 850 (15.95) | 1375 (17.8) | 499 (10.09) | 553 (10.64) | 1626 (27.36) | 1887 (33.01) |
| 11-15 | 630 (11.85) | 531 (10.34) | 622 (11.02) | 495 (8.95) | 170 (3.19) | 354 (4.58) | 100 (2.02) | 143 (2.75) | 390 (6.56) | 479 (8.38) |
| 16-20 | 159 (2.99) | 142 (2.77) | 187 (3.31) | 134 (2.42) | 60 (1.13) | 85 (1.1) | 32 (0.65) | 51 (0.98) | 111 (1.87) | 114 (1.99) |
| 21-30 | 112 (2.11) | 69 (1.34) | 134 (2.37) | 73 (1.32) | 34 (0.64) | 57 (0.74) | 20 (0.4) | 45 (0.87) | 79 (1.33) | 49 (0.86) |
| 31-100 | 66 (1.24) | 54 (1.05) | 64 (1.13) | 46 (0.83) | 33 (0.62) | 68 (0.88) | 24 (0.49) | 47 (0.9) | 54 (0.91) | 8 (0.14) |
| 101-200 | 9 (0.17) | 1 (0.02) | 1 (0.02) | 1 (0.02) | 3 (0.06) | 6 (0.08) | 3 (0.06) | 5 (0.1) | 3 (0.05) | 0 (0.0) |
| 201-300 | 2 (0.04) | 0 (0.0) | 0(0.00) | 0(0.0) | 1 (0.02) | 1(0.01) | 0 (0.0) | 0 (0.0) | 1 (0.02) | 0 (0.0) |
| 301-400 | 0 (0.0) | 0 (0.0) | 0 (0.0) | 0 (0.0) | 0 (0.0) | 0 (0.0) | 0 (0.0) | 0 (0.0) | 1 (0.02) | 0 (0.0) |
| 401-500 | 0 (0.0) | 0 (0.0) | 0 (0.0) | 0 (0.0) | 0 (0.0) | 0 (0.0) | 0 (0.0) | 1 (0.02) | 0 (0.0) | 0 (0.0) |
| Total | 5318 | 5134 | 5644 | 5530 | 5328 | 7724 | 4945 | 5199 | 5942 | 5716 |

### Most *P. falciparum* proteins with long TRs are exported

*Plasmodium falciparum* is the most virulent malaria causing species that is responsible for most disease associated fatalities [66]. The virulence of Pf is largely attributed to its ability to cytoadhere in the host microvasculature *via* parasite proteins exported to the infected red blood cell surface [67], [41]. Therefore, we analysed the TR regions of Pf (reference strain 3D7) for their length and coverage in an attempt to understand the contribution of repetitive sequences in Pf biology. The percentage coverage of the repeat region in these proteins ranged from 0.34% (PF3D7_0630300 encoding for DNA polymerase epsilon catalytic subunit A) to 99.7% (PF3D7_1211800 encoding for polyubiquitin - a component of polyubiquitin ligase) respectively. Some of the other Pf proteins with high occupancy of TR regions are Pf3D7_1038400 (gametocyte specific protein with 89.25% coverage), Pf3D7_0501400 (FIRA - interspersed repeat antigen with 84.43% coverage), PF3D7_1035200 (S antigen with 76.58% coverage) and PF3D7_0201900 (erythrocyte membrane protein 3 with 74.59% coverage). While polyubiquitin is a universal protein that carries the ubiquitin domain placed in tandem 5-6 times in all eukaryotes, the other proteins with high TR coverage are specific to *Plasmodium*. Several of the above proteins are exported outside the parasite confines and localize to various subcellular positions like the iRBC surface, iRBC cytoplasm, Maurer’s clefts, extracellular vesicles, symbiont containing vacuoles, knobs *etc*. We also assessed the cellular localization and annotations of all proteins with more than 50% TR coverage (Table S1). Several of these proteins belong to the Pf exportome with or without the occurrence of *Plasmodium* export element (PEXEL) motif. PEXEL motif (consensus sequence RxLxE/D/Q) is a pentameric sequence whose presence at the N terminus of proteins commits them to be exported from the parasite into the host erythrocyte where it resides [68].

### TRs of *P. falciparum* are enriched in Asparagine and acidic amino acids

Repeat regions in eukaryotes are often enriched in polar uncharged amino acids like Gln, Asn, Ser, Pro and Thr along with acidic amino acids (Glu & Asp) and small amino acids Gly and Ala [69], [70]. We have analysed the amino acid composition of TR regions of *Plasmodium falciparum* for comparison with other eukaryotes. Tandem repeats in Pf were particularly rich in Asn, Lys and acidic amino acids Asp and Glu (Figure 2a). Asn was present most frequently (30.47%) followed by Asp (13.21%), Lys (10%) and Glu (9.68%). The percentage of Trp was the lowest (0.02%).

**Figure 2:**
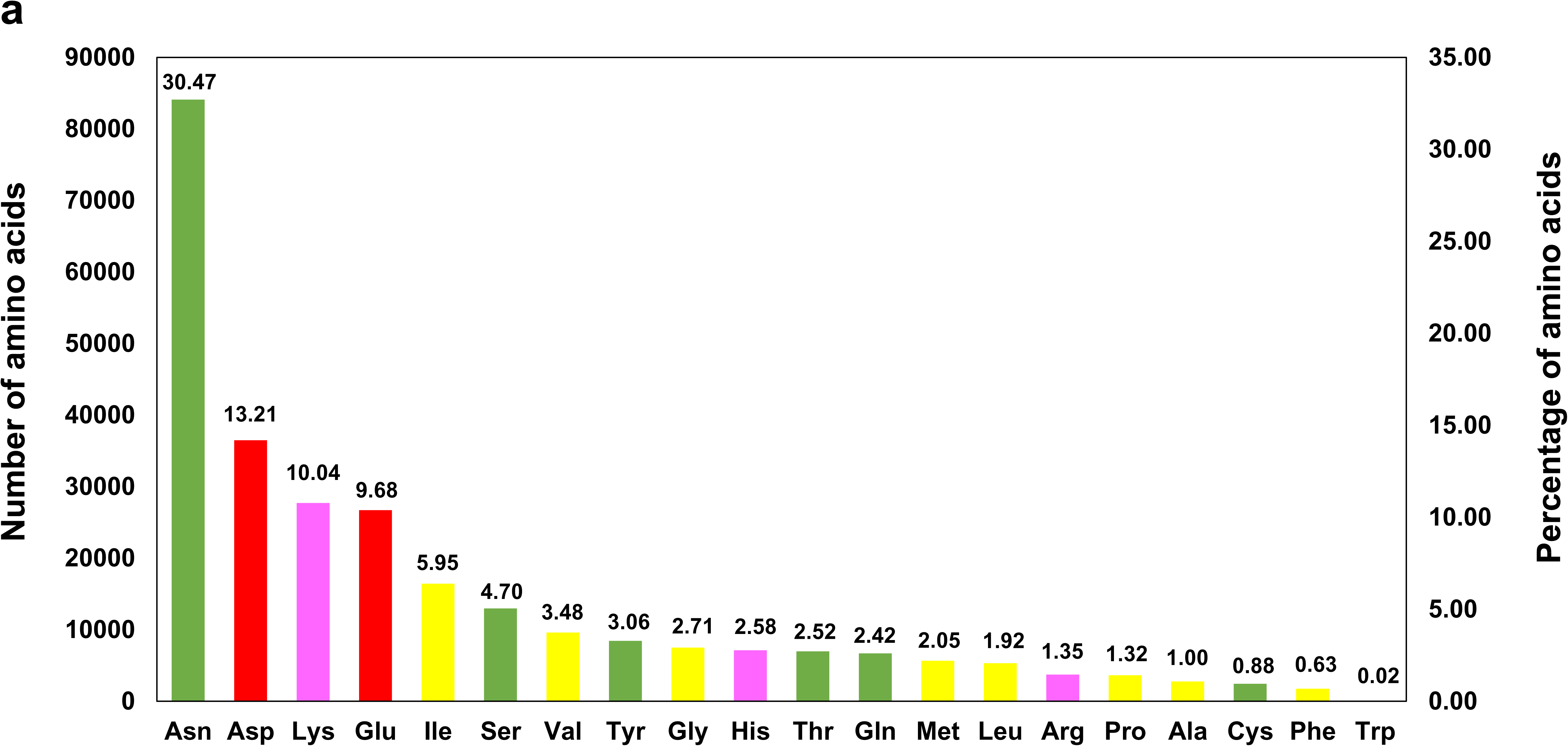
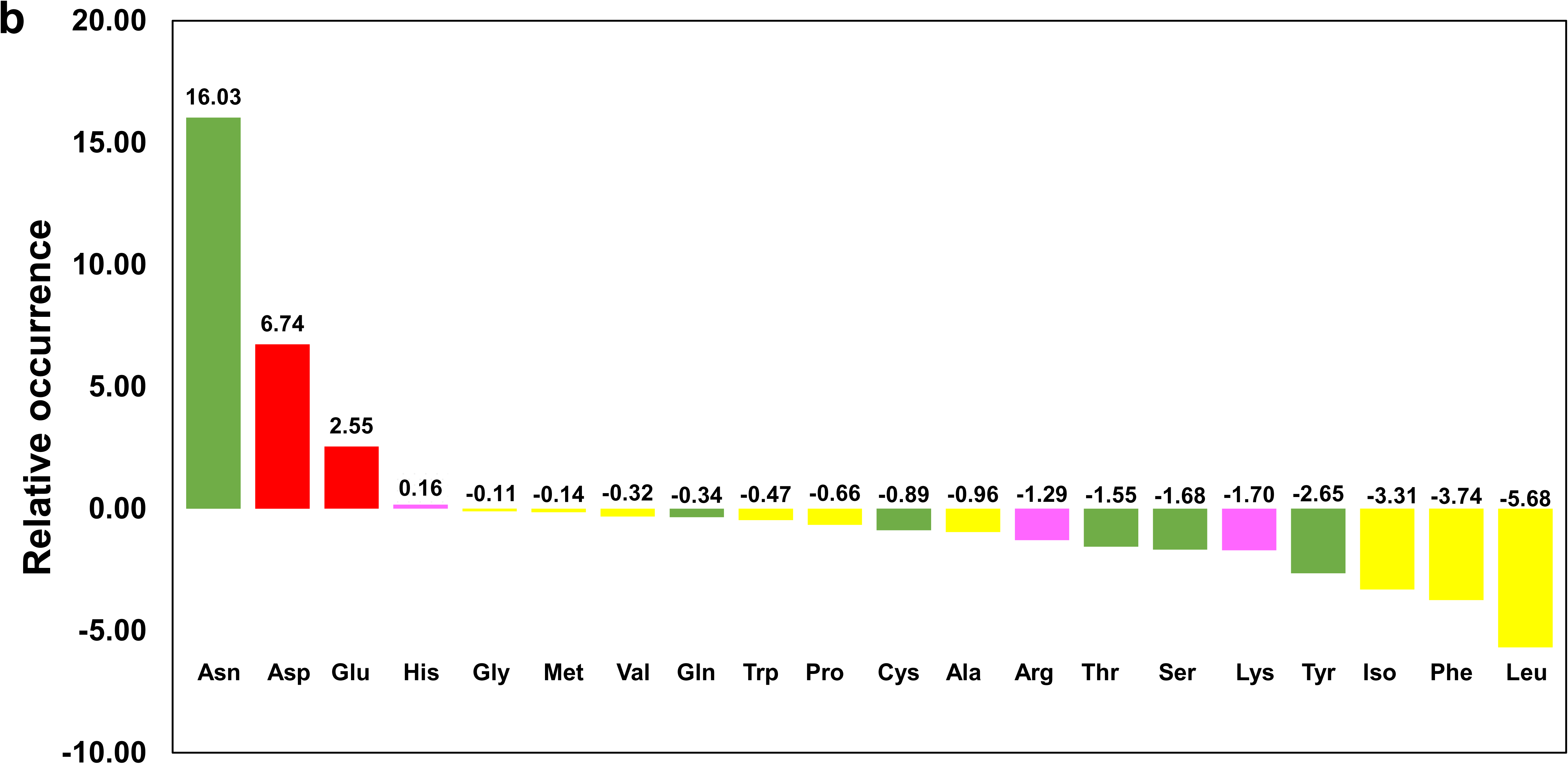
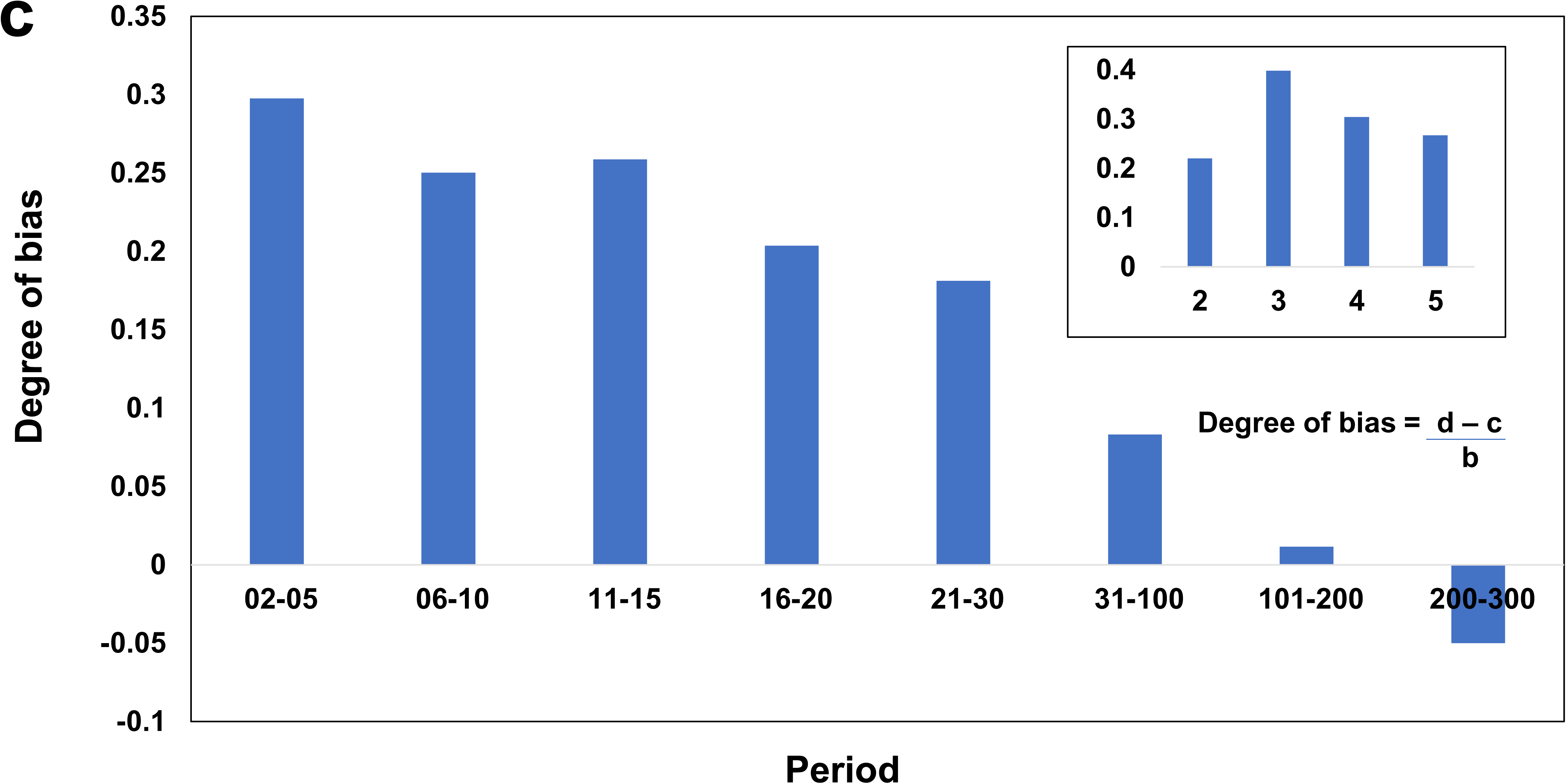
Amino acid usage in P. falciparum repetome. a) Bar diagram representing the amino acid composition (number and percentage) of each standard amino acid in the repeat content of Pf. X-axis depicts each amino acid as labelled and Y axis shows the number and percentage as labelled. Percentage of each amino acid is also inscribed above the respective bar, and was calculated considering total number of amino acids in the repetome as 100%. The bars for different amino acids are color coded according to their character *viz.* green: polar, red: acidic, pink: basic and yellow: non-polar. b) Relative occurrence (RO) of standard amino acids i.e. difference between percentage of an amino acid in repetome and percentage of that amino acid in total proteome. Amino acids are denoted by their single letter code. RO values are inscribed above each bar. Positive values indicate over-representation; negative values indicate under-representation. c) Chart representing Asn usage bias in Pf repetome (in comparison with total proteome) for the periods 2-5, 6-10, 11-15, 16-20, 21-30, 31-100, 101-200 & 201-300. Formula for the calculation of Asn usage bias is shown on the chart, where b = Total number of amino acids in the TR region of a given period; c = Expected number of Asns in each period in the absence of bias and d = Number of Asns present in each period. Usage bias for periods 2, 3, 4 & 5 is shown in figure inset.

We also assessed the relative occurrence of each standard amino acid in the repeat regions versus the total Pf proteome (percentage of an amino acid in repetome minus percentage of that amino acid in total proteome) (Figure 2b). While Asn (+16.03%), Asp (+6.74%), Glu (+2.55%) and His (+0.16%) were found to be overrepresented in the TR regions, all the other amino acids were underrepresented. Non polar aliphatic amino acids Leu (-5.68%) & Ile (-3.31%) and the aromatic amino acids Phe (-3.74%) & Tyr (-2.65%) were among the most underrepresented residues. The contribution of Asn and Asp in the repeat regions was found to be more than double their percentage in the total Pf proteome (Asn: 14.44% and Asp: 6.47%). While the occurrence of Glu was only marginally higher in TRs as compared with the total proteome (7.13%), Lys was more poorly represented in repeat regions in comparison with the Pf proteome (11.74%) (Table S2). Pf fulfills its requirement of amino acids largely through hemoglobin degradation and import from the host cell [71]. Although amino acids obtained from external sources are enough for parasite survival, its genome still encodes for enzymes that synthesize Asn, Asp, Glu, Gln, Gly and Pro. Interestingly, Pf shows an overrepresentation of three of the amino acids it synthesizes i.e. Asn, Asp and Glu in its repeat content.

### Short and intermediate TRs in Pf are biased towards Asparagine usage

Asn is the most abundant amino acid in *Plasmodium falciparum* proteome, which expresses the enzyme Asparagine synthase for production of this amino acid [71]. Asn synthase gene knockout Pb parasites showed delayed development of liver and intra erythrocytic stages [72]. Formation of ookinetes, oocysts and sporozoites in the mosquito stages of the mutant parasites was also significantly reduced. It has been earlier reported that homorepeats in Pf are dominated by Asn which was believed to result from AT richness (81% AT content) of the Pf genome [21]. This hypothesis was later negated since Asn (14.44%) occurred much more commonly than Lys (11.74%) in the Pf proteome despite sharing an equivalent AT content in their codons. Also, Asn occurrence in Pv and Pk matches that of Pf despite the relative GC richness of their genomes (Pv: 42% GC and Pk: 38% GC content respectively) [73], [74]. It has been proposed that Asn homorepeats may be subject to positive selection in *Plasmodial* species (54,). Keeping the above in mind, we have analysed the Asn content of TRs in the Pf proteome as a function of their period. The degree of bias in Asn usage was determined for the periods 2-5, 6-10, 11-15, 16-20, 21-30, 31-100, 101-200 and 201-300 as explained in the Methods section of this article (Figure 2c). Asn usage showed the greatest bias in short 2-5 mer TRs (0.297). Among the 2-5 residue long TRs, 3 mers showed the strongest bias (0.398) followed by 4 mers (0.304), 5 mers (0.267) and then 2 mers (0.221) (Figure 2c inset). TRs with periods 6-10 and 11-15 showed nearly equal Asn usage bias (0.25 and 0.258 respectively) which was relatively lower than the period 2-5. As the length of period increased further, there was a continuous decline in the bias for usage of Asn. In order to understand which of the periods showed a significant Asn usage bias, we calculated the Gly usage bias in the entire proteome (0.0153) for comparison. Gly was chosen since this amino acid had shown the least difference in its relative occurrence in the repetome vs total Pf proteome (-0.01%). Asn usage bias seemed to be insignificant in periods larger than 100 as these values were less than double of Gly usage bias. Higher Asn usage bias in shorter TRs may be a result of slippage events at the time of DNA replication. It seems logical that Asn usage in longer repeats is similar to the total Pf proteome since such stretches are formed by duplication events, and not DNA replication errors.

### Gene ontology analysis of TR containing proteins of *P. falciparum*

Identification of TRs in Pf by using XSTREAM (copy number 2, period 3, minimum TR domain length 20) yielded 1598 proteins (Supplementary file 2). Of these, two were annotated as hypothetical proteins and 511 had no known function. The GO terms listed against cellular component, molecular function and biological processes for each of these proteins were run through TB tools in an attempt to obtain the number of annotated proteins corresponding to each GO class. The total number of proteins found to have annotations in different GO classes were 1038 under the cellular component group, 658 under molecular functions and 626 under biological processes.

#### Cellular component

The 1038 proteins obtained upon GO analysis for cellular components were sub-classified manually as per their association with the membrane, nucleus, host cell, specific organelles and / or other macromolecules to form complexes (Figure 3a). The maximum number of TR containing proteins (369) were annotated to localize in the cytoplasm, followed by the nucleus (282), then as part of protein - containing complexes (182) or belonging to the host cellular component (126). 83,79 and 66 proteins were found to be present in the apicoplast, membranes and Maurer’s clefts respectively. It is pertinent to mention here that one protein can be present in multiple sub-classes (cellular localization) and therefore the number of proteins present in each category would not add up to the total number in a specific GO class.

**Figure 3:**
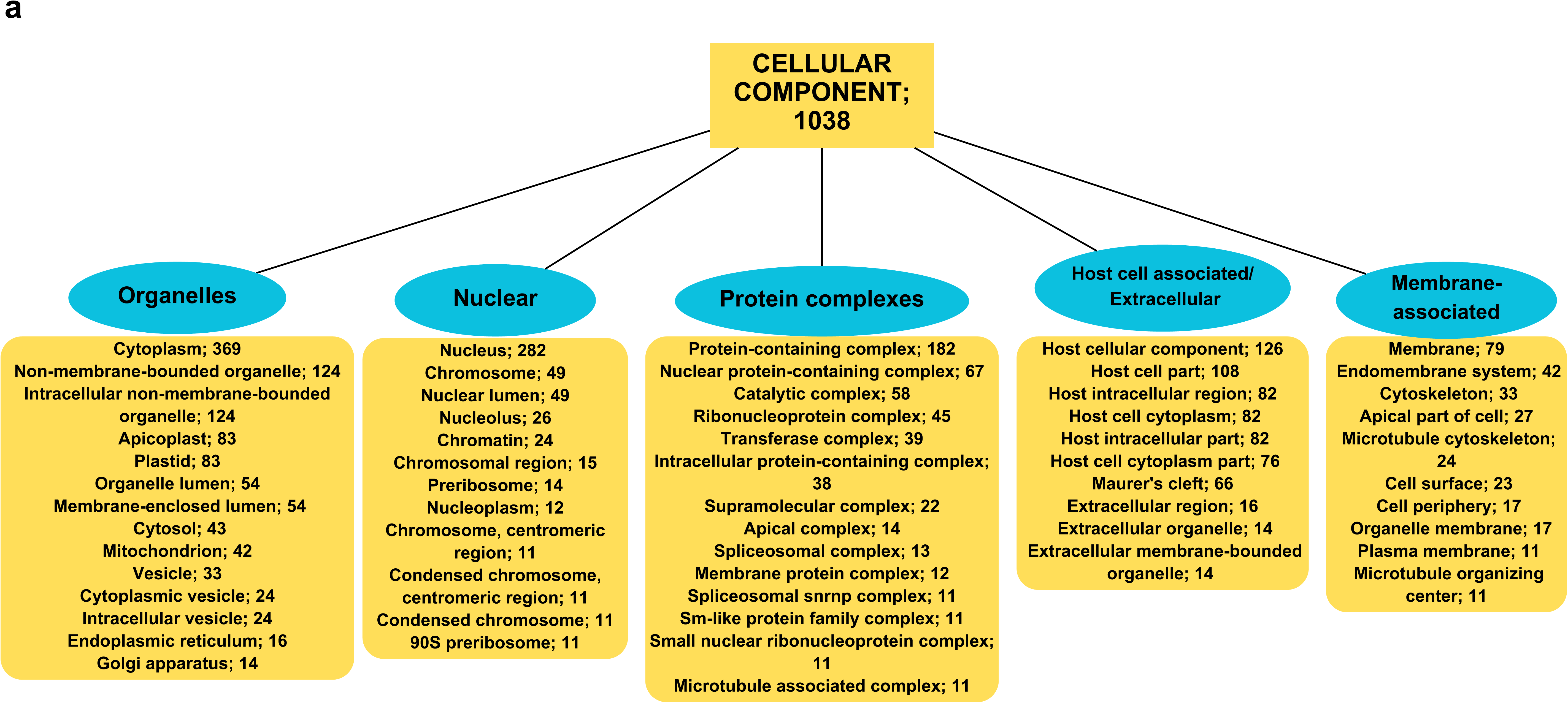
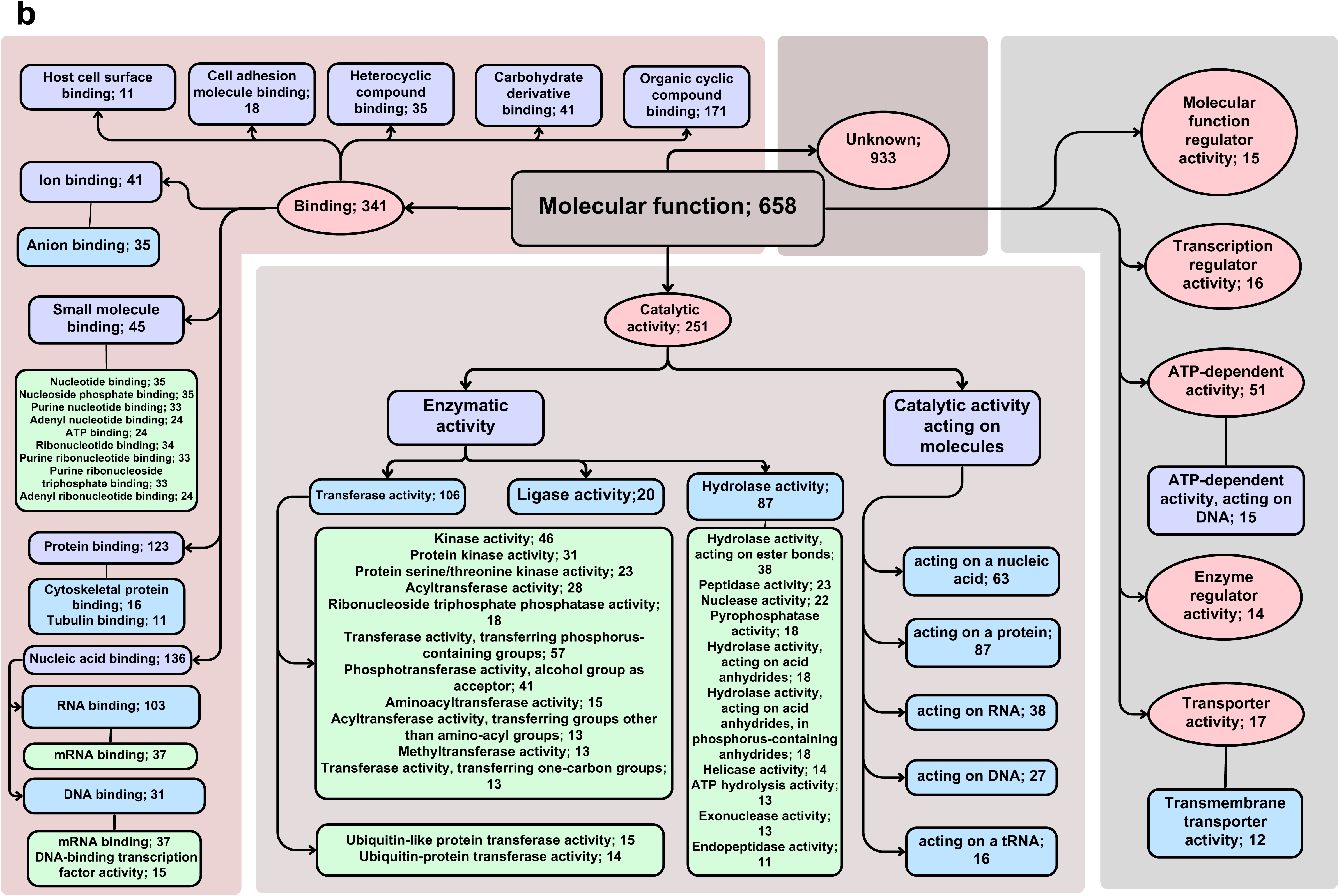
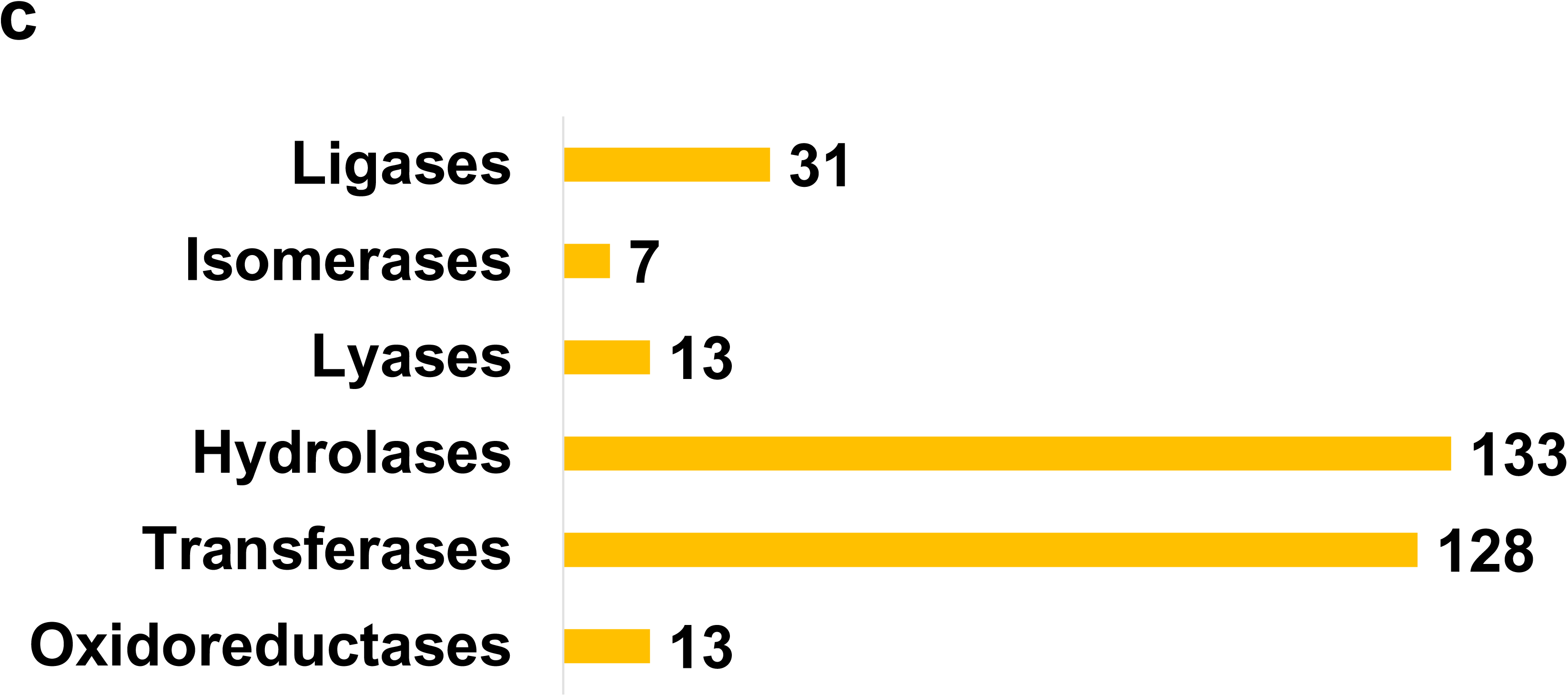
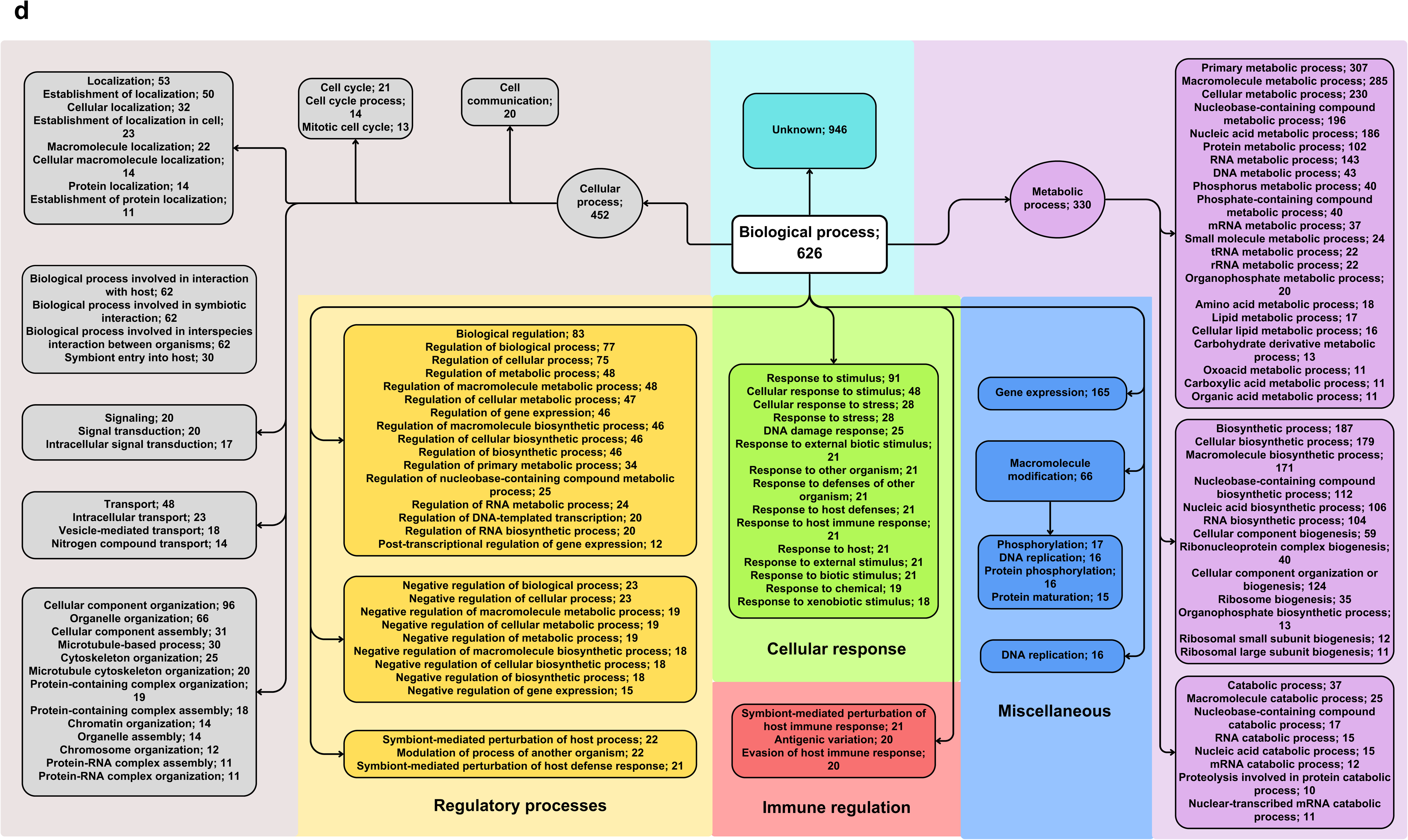
Gene ontology analysis of Pf repetome. 1572 proteins of Pf repetome identified by XSTREAM (copy number 2, period 3, minimum TR domain length 20) classified according to a) GO terms (Cellular component) b) GO terms (Molecular function) c) Enzyme commission numbers (Enzyme classes) d) GO terms (Biological processes).

#### Molecular function

Proteins of the second GO class describing molecular functions (658 annotated) were grouped into seven categories based on the activities they perform *viz.* binding (341), catalytic (251), ATP dependent activity (51), transporters (17), transcription regulators (16), molecular function regulators (15) and enzyme regulators (14) (Figure 3b). Proteins with binding activity were annotated to bind diverse molecules like organic cyclic compounds (171), nucleic acids (136), proteins (123), small molecules binding e.g. nucleotide binding proteins (45) and carbohydrate derivatives & ions binding (41 each). Proteins carrying catalytic activity had been further classified based on known enzyme classes or the molecules they act upon. Interestingly, TR containing proteins belonged to only three enzyme classes *viz.* transferases (106), hydrolases (87) and ligases (20) with the ability to act on proteins and/ or nucleic acids.

As an alternate approach to understanding the enzymatic potential of TR containing proteins (1572 proteins), we identified and listed the ones associated with EC (enzyme commission) numbers. This yielded 325 proteins that were sorted and classified based on their EC classes. This number was much larger than the number of TR containing proteins identified by GO analysis to carry enzymatic activity. Here, the number of proteins in hydrolase class (133) superseded transferases (128). Ligases, oxidoreductases, lyases and isomerases carried 31, 13, 13 and 7 proteins respectively (Figure 3c). Most of the TR containing transferases were classified as non-specific serine/threonine protein kinases, and most hydrolases had RNA helicase (21), DNA helicase (14) or Protein-serine/threonine phosphatase (11) activities (Table 3).

**Table 3:** TR-containing enzymes. Table shows the distribution of TR-containing proteins based on their enzyme commission (EC) numbers (class and sub-class).

| Enzyme class | Enzyme sub-class | No. of proteins |
| --- | --- | --- |
| Transferases | Non-specific serine/ threonine protein kinase | 31 |
| Hydrolases | RNA helicase | 21 |
| Hydrolases | DNA helicase | 14 |
| Hydrolases | Protein-serine/ threonine phosphatase | 11 |
| Transferases | Methyltransferases | 8 |
| Transferases | Histone acetyltransferase | 7 |
| Hydrolases | Ubiquitinyl hydrolase 1 | 7 |
| Transferases | DNA-directed RNA polymerase | 6 |
| Transferases | DNA-directed DNA polymerase | 6 |
| Hydrolases | Cysteine endopeptidases | 5 |
| Ligases | Long-chain-fatty-acid-CoA ligase | 5 |
| Transferases | RING-type E3 ubiquitin transferase | 4 |
| Transferases | Non-specific protein-tyrosine kinase | 4 |
| Transferases | Cyclin-dependent kinase | 4 |
| Lyases | Pseudouridylate synthase | 4 |
| Transferases | 1-phosphatidylinositol-4-phosphate 5-kinase | 3 |
| Hydrolases | Lysophospholipase | 3 |
| Lyases | tRNA pseudouridine (38-40) synthase | 3 |

#### Biological processes

The third GO classification on the basis of biological processes was used to classify TR containing proteins on the basis of metabolic, cellular and regulatory processes they are involved in. Proteins associated with cellular response, immune regulation, gene expression, macromolecule modification and DNA replication were also included. The largest number of proteins were categorized to be part of cellular processes (452) (Figure 3d), followed by metabolic processes (330), gene expression (165), response to stimulus (91) and biological regulation (83) etc. While the metabolic processes included both catabolic and biosynthetic processes, the cellular processes category consisted of proteins with roles in structural organization of cellular/ subcellular components, biological interactions, cellular localization, transport, cell cycle, signaling and cell communication.

### Pf Variant surface antigen (VSA) families have low repetitive content

Short tandemly repeated sequences in Pf proteins have been related to generation of antibody responses against field isolates [56]. Surface antigens of the malaria parasite are considered as important targets of host antibody response, which forms a component of naturally acquired immunity in patients [75]. *Plasmodium falciparum* (strain 3D7) hosts a total of five variant surface antigen (VSA) families *viz.* Pf erythrocyte membrane protein 1 (PfEMP1; 65 proteins), sub-telomeric open reading frame (STEVOR; 32 proteins), surface-associated interspersed gene family (SURFIN; 7 proteins), repetitive interspersed family (RIFIN; 157 proteins), and Pf Maurer’s cleft two transmembrane (PfMC-2TM; 11 proteins) [76]. Proteins expressed by all these sub-telomeric multigene families except PfMC-2TM localize to the cell surface. Members of these families are involved in generation of antigenic diversity, immune evasion, merozoite invasion, rosetting and cytoadherence. Considering the antigenic nature of VSAs, we have assessed the repeat content of Pf VSAs to understand whether their antigenicity is related to the TR sequences they carry.

71.4% of the SURFINs (5 out of 7), 32.3% of PfEMP1s (21 out of 65) and only 0.63% of RIFINs (1 out of 157) were found to contain tandem repeats (XSTREAM search parameters: minimum copy number 2, minimum period 3 and minimum TR region length 20). None of the STEVOR or PfMC-2TM proteins contained repeat sequences. Raghavan *et. al.* had identified antigenic peptides from 1648 seroreactive proteins (∼30% of Pf proteome) from VSA families and other antigens expressed at different stages of the parasite life cycle [56]. The antibody response to antigens with repetitive content was shown to be relatively short lived and reliant upon pathogen exposure when compared with response to non-repetitive antigens. We compared the identified TR containing VSAs with the reported seropositive peptide data that revealed all proteins with repeats to be seroreactive, particularly in a high disease transmission setting. Also, several of the repeat regions were found to lie within seroreactive peptides (Table 4). Repetition of epitopes increases their valency and holds the ability to alter the nature of the host immune response [77].

**Table 4:**
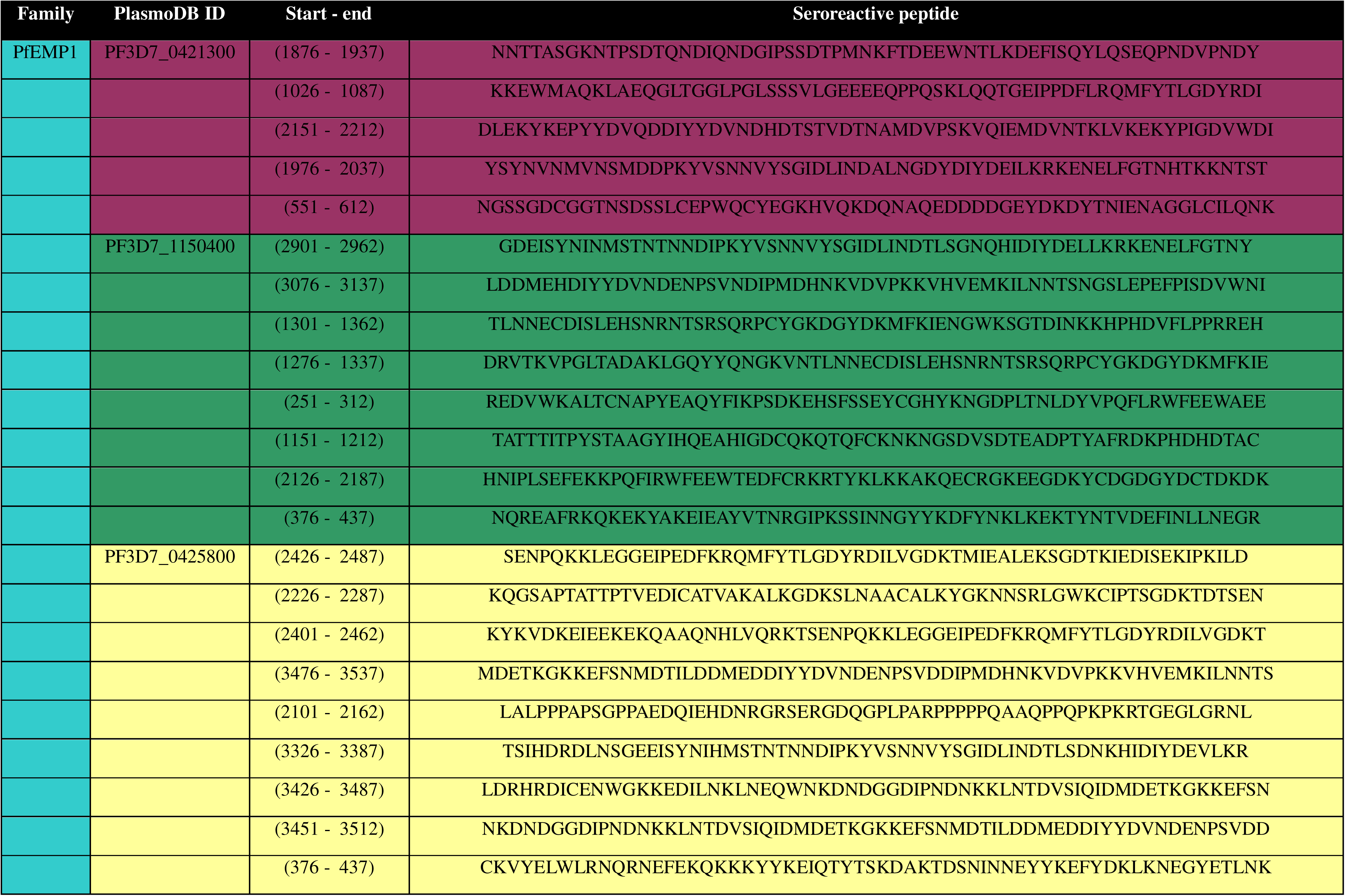
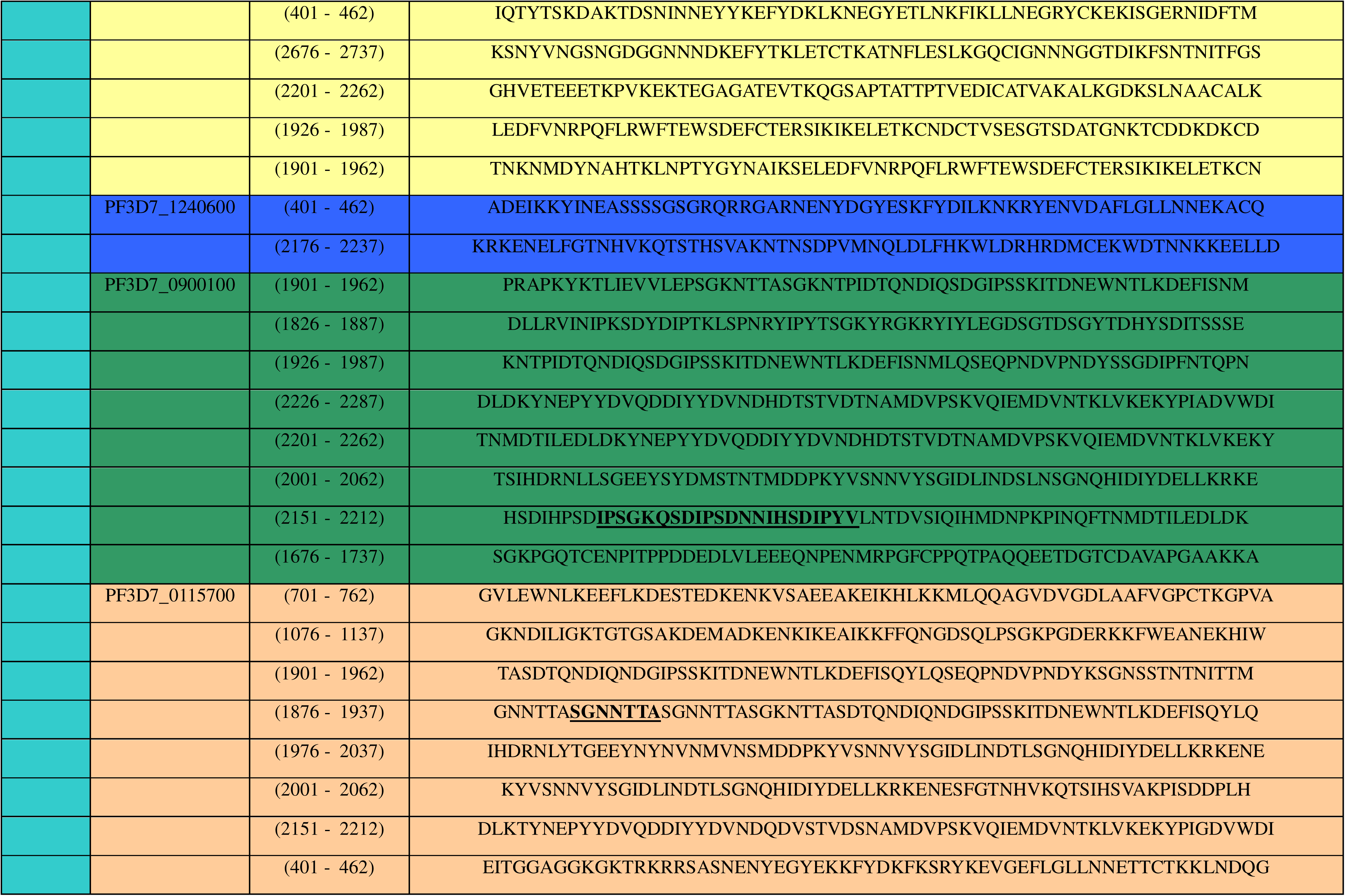
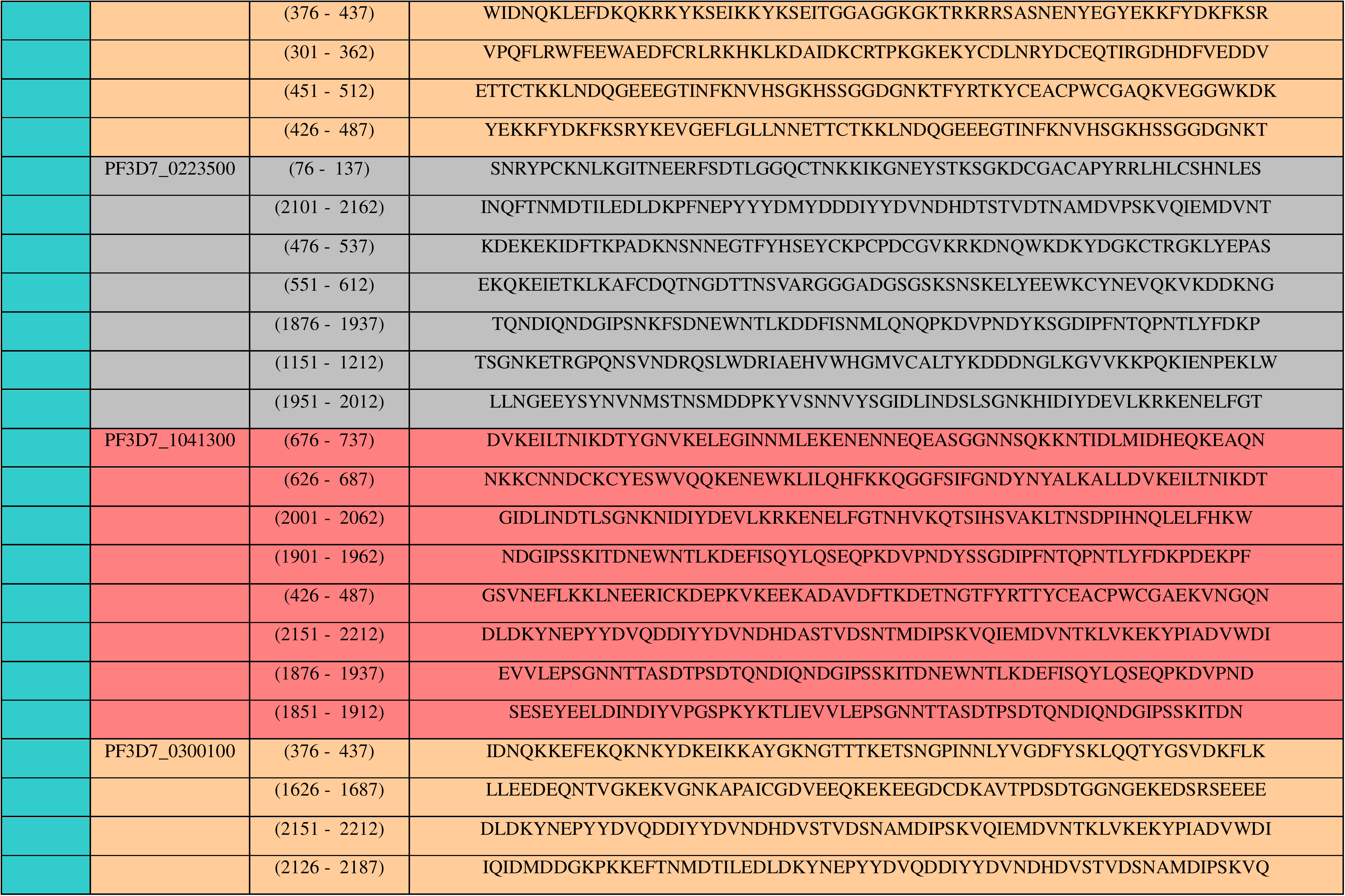
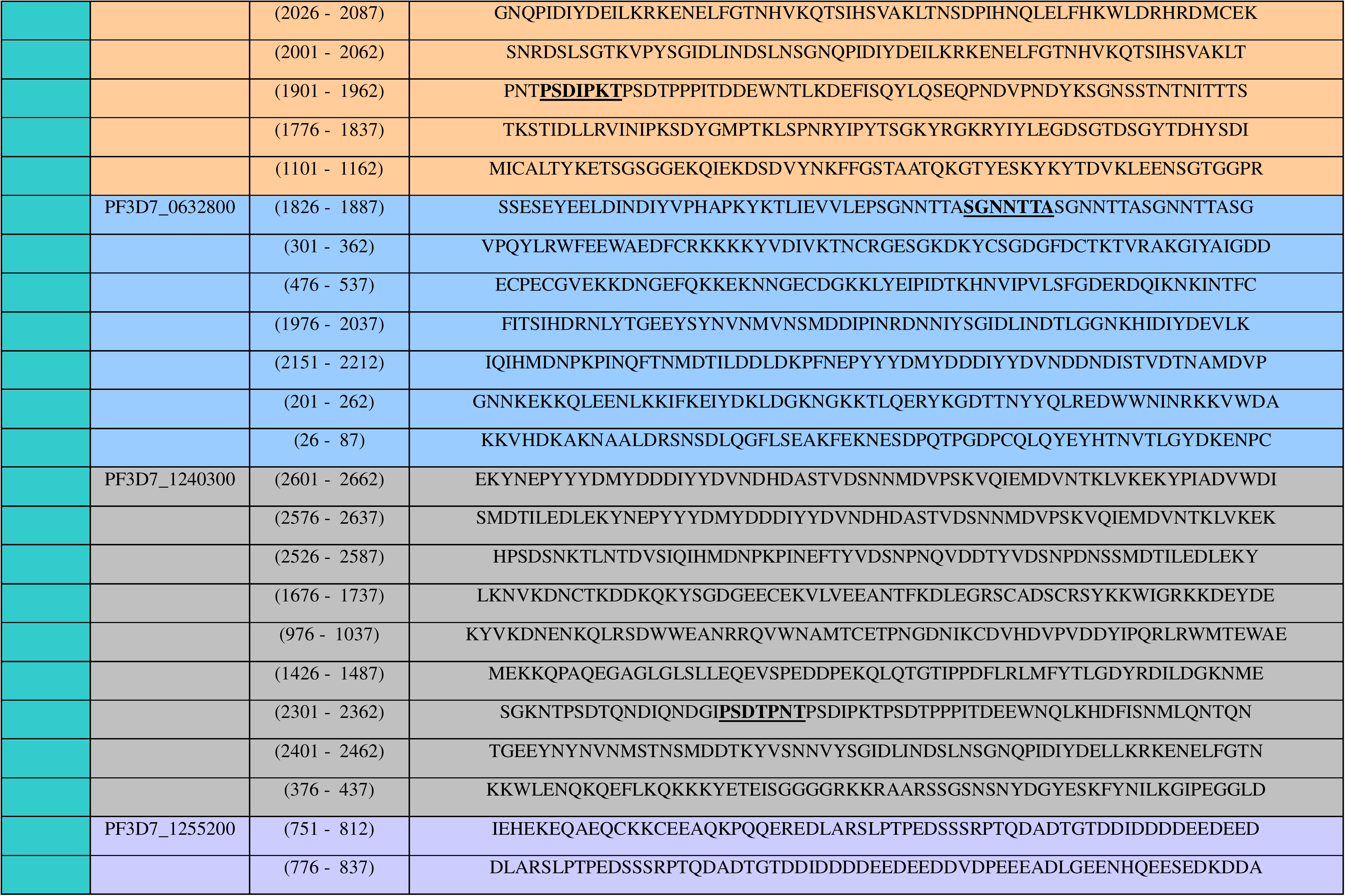
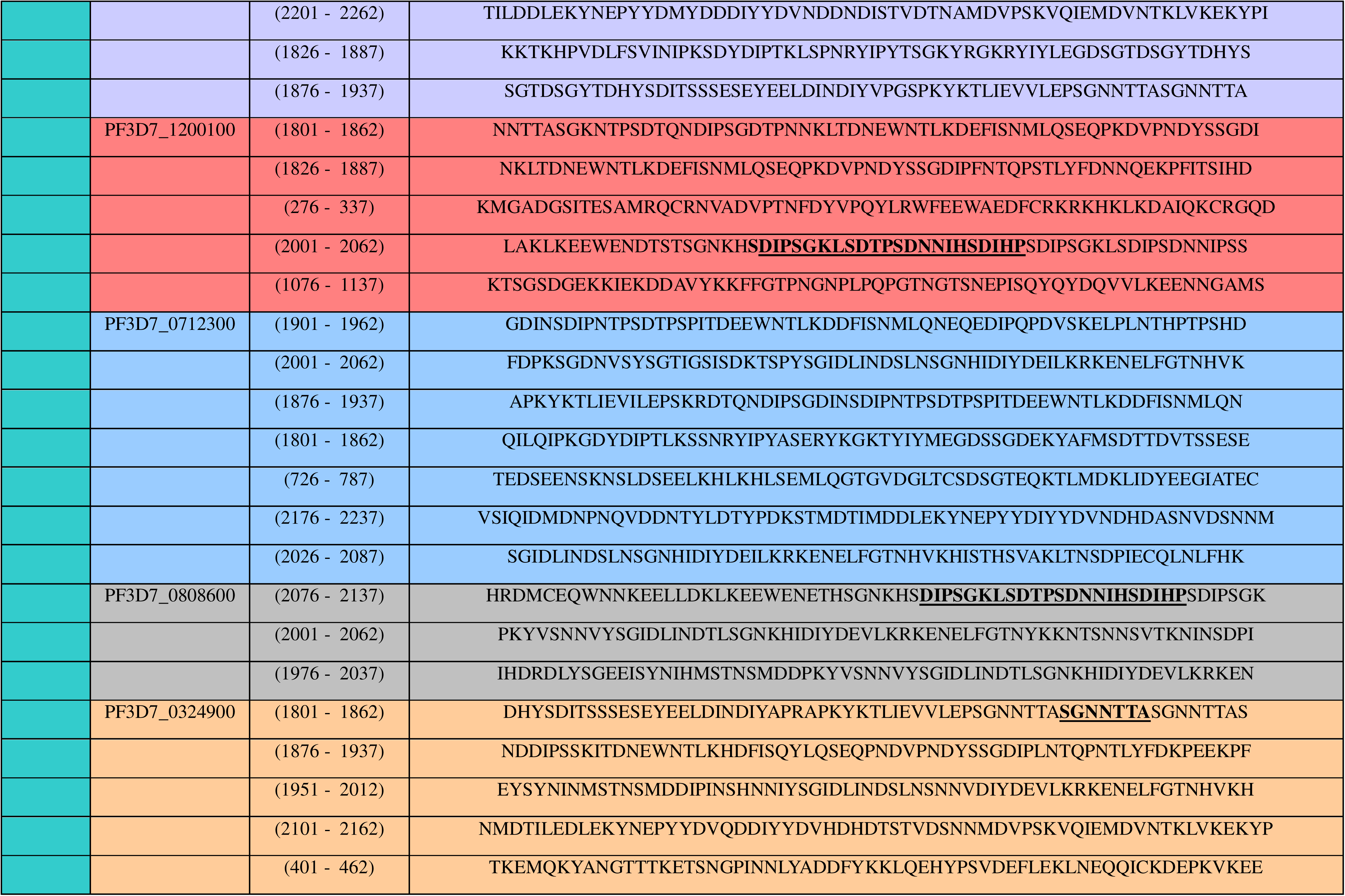
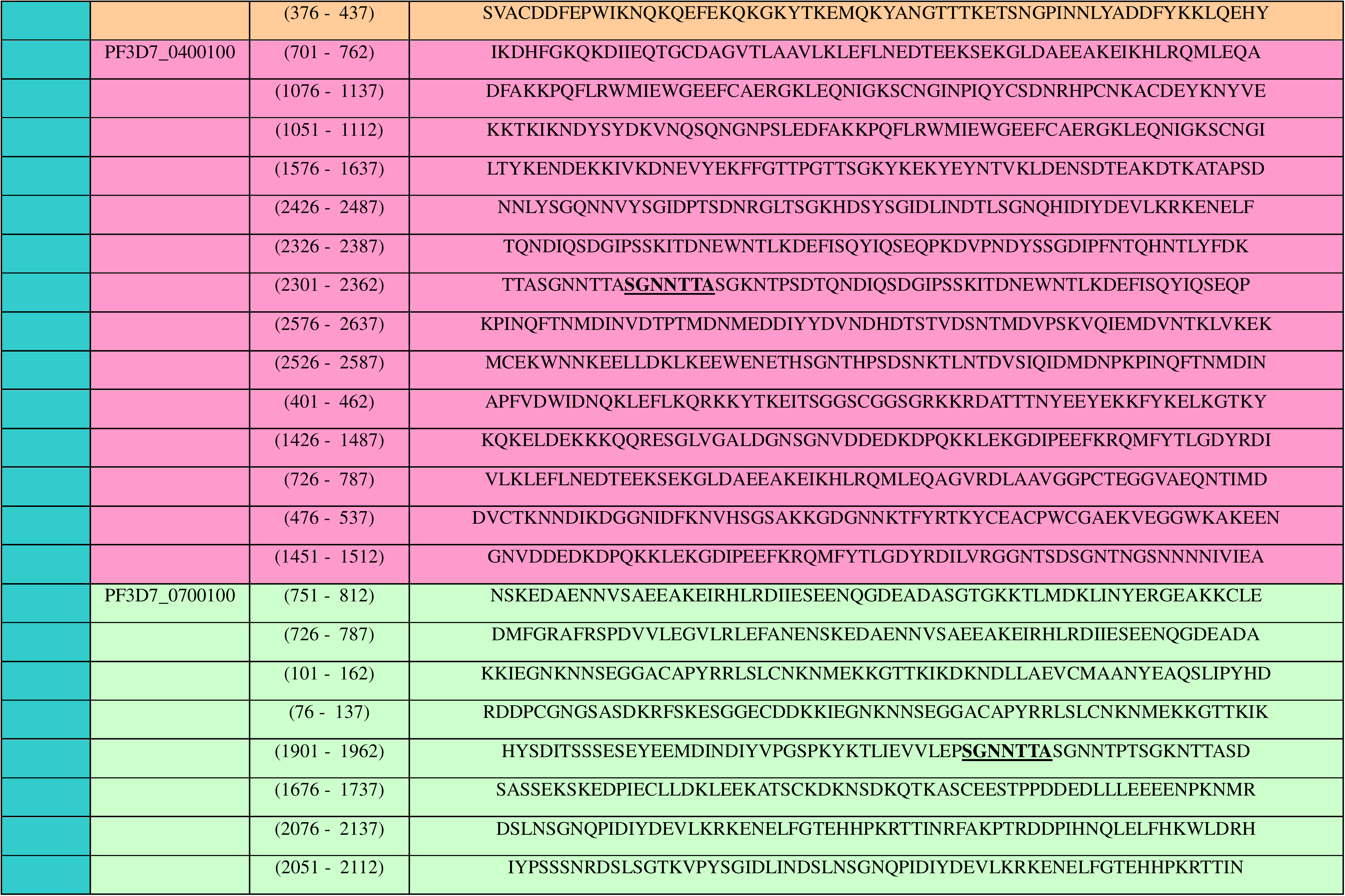
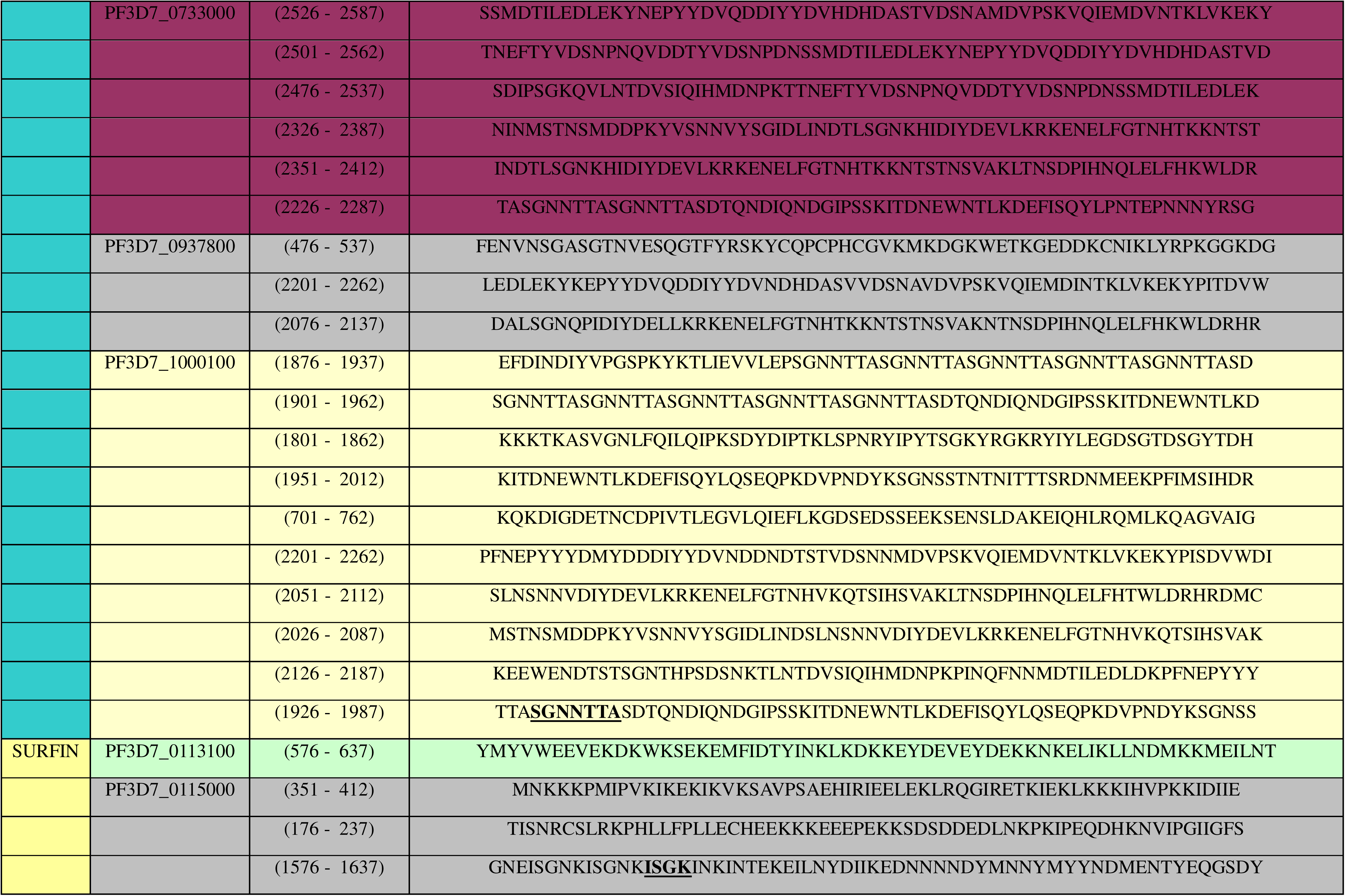
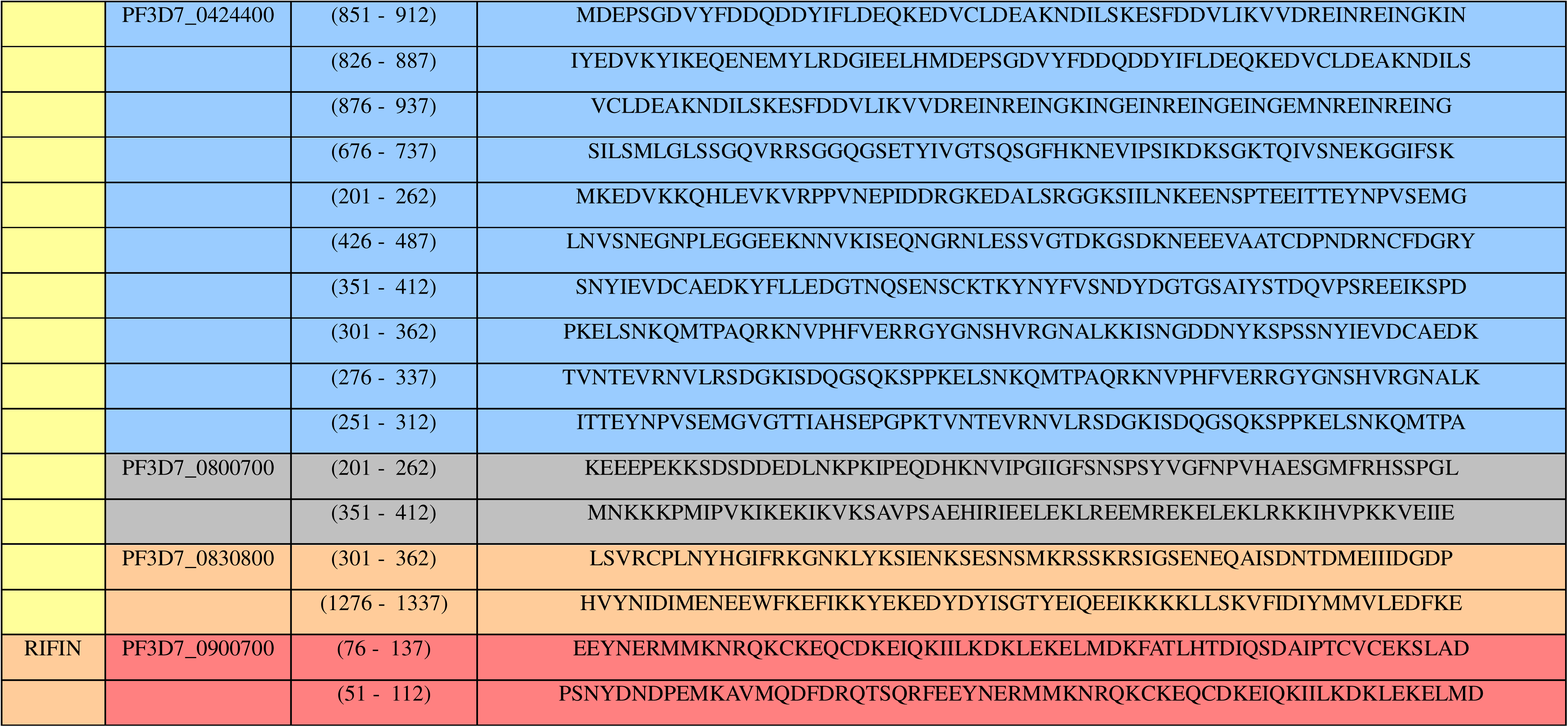
Seroreactive peptides in TR containing proteins of Pf variable surface antigen (VSA) families. Table lists TR-containing proteins of RIFIN, SURFIN and PfEMP1 families, along with sequences of their seroreactive peptides as reported by Raghavan *et. al.* PlasmoDB ID, start point, end point and sequences of respective seroreactive peptides are mentioned. TR units present within seroreactive peptides are highlighted in bold.

The only RIFIN carrying a TR sequence was ‘PF3D7_0900700’, an A-type RIFIN that contained an imperfect 10 mer sequence repeating only twice (AAKKLAAEAG AAKGLAAGAE). Its repeat region was present almost in the middle of its VSA_Rifin domain. The SURFINs that carried oligomeric TRs were 1.1 (PF3D7_0113100), 1.3 (PF3D7_0115000), 4.2 (PF3D7_0424400), 8.2 (PF3D7_0830800) and 8.3 (PF3D7_0800700). None of the SURFINs had appreciable TR region coverage as compared with the entire protein (maximum coverage 0.04% for SURFIN 1.1). SURFINs 8.2 and 4.2 carried two repetitive stretches each, while the others had only one. SURFIN 4.2 is reported to be a component of the SURGE complex where it interacts with RON4 (rhoptry neck protein 4) and GLURP (Glutamate rich protein) to facilitate merozoite invasion [78], [79], [80]. Its intracellular tryptophan rich domain (WRD) is involved in transport from Maurer’s clefts to the iRBC membrane. Both the TRs it carries lie within its WRD. Other SURFINs carrying TR regions have not been functionally characterized till date. SURFIN 1.1 has been reported to be highly immunogenic as it experiences a strong diversifying immune selection [81]. The repeat regions of SURFINs 1.3 and 4.2 lie within the seroreactive peptides determined by Raghavan *et. al*.

### Repeat content of intracellular domains of PfEMP1 is seroreactive

The antigenically variant PfEMP1 proteins are encoded by *var* genes (∼ 60 in number) that have a two-exon structure [82], [83]. The first exon encodes their ectodomain comprising multiple duffy binding like (DBL) domains and cysteine-rich inter domain regions (CIDR) among others, and engages a variety of host receptors resulting in cytoadherence of parasitized erythrocytes in the microvasculature to effect malaria related complications. The second exon codes for the intracellular acidic terminal segment (ATS) that resides in the iRBC cytosol, and interacts with both host and parasite expressed proteins to form a protein meshwork [84], [85]. PfEMP1s display antigenic switching where only one member of the family expresses itself at a given time [83]. Our results show that out of the twenty-one PfEMP1 proteins carrying TRs, repeats of fifteen were present only on their ATS domains (Table S3). Of these, PF3D7_0632800 had three TR regions with consensus sequences PSDTPKT, SDIPSGKL and SGNNTTA (copy numbers ∼3, 3 and 5 respectively) spread throughout its ATS. Interestingly, the 7 mer ‘SGNNTTA’ was also found in ATS domains of 7 other PfEMP1 proteins (PF3D7_0324900, PF3D7_0400100, PF3D7_0700100, PF3D7_0733000, PF3D7_0937800, PF3D7_1000100 & PF3D7_1255200) repeating 3-5 times. PF3D7_0733000 had two TR stretches in its ATS region with SGNNTTA repeating 4.14 times and an 11 mer ‘SDIHPSDNNIH’ repeating twice. The longest repeat in ATS was the 23 mer consensus sequence ‘SDIPSGKLSDIPSDNNIHSDIHP’ present in PF3D7_1200100 repeating twice. A similar sequence was also present in the ATS domains of two other PfEMP1s *viz.* PF3D7_0808600 and PF3D7_0900100. Each of these consensuses repeat sequences were found to be present within seroreactive peptides from multiple PfEMP1 proteins (Table 4). Such shared motifs are a common feature of the PfEMP1 family that may act as cross-reactive epitopes reducing the effectiveness of the host immune response to the malaria parasite [56].

The ectodomains of a total of six PfEMP1 members contained tandem repeats that were invariably present in the linker regions between two domains. TR units of three such PfEMP1 proteins were exceptionally rich in acidic amino acids (PF3D7_1150400, PF3D7_0421300 & PF3D7_0425800) (Table S3). Ectodomains of PF3D7_1150400, PF3D7_1255200 and PF3D7_0223500 had TR regions with high proline and glutamine content. Highly seroreactive peptide stretches of Pf are reported to carry both acidic amino acids Asp & Glu, and basic amino acids Lys & Arg [56]. PF3D7_1000100 and PF3D7_1255200 carried one repeat region each in their ecto and ATS domains. None of the repeat motifs of the ectodomains could be found in the list of seroreactive peptides. Our results corroborate the finding that ATS domains of PfEMP1s are more seroreactive than their ectodomains. This possibly results from their sequence conservation across PfEMP1 variants leading to repeated display of B cell epitopes of ATS to the host immune system despite antigenic switching [56].

### Repeat regions of several Pf proteins are structured

Low complexity TR regions can either be unstructured or may adopt a structure [86]. Here, we have studied structures of a total of 68 TR regions from a variety of Pf proteins that were representative of each available period (ranging from 4 to 250), and had the highest copy number (Table 5). Solved structures for none of these repeat units were available in the Protein data bank (PDB) [57]. Predicted structures of TR regions of these proteins were obtained through AlphaFold [58], [59]. AlphaFold is an artificial intelligence generated database containing three-dimensional protein structures predicted from their primary sequence. Of the 68 studied proteins, structures of 43 were available on AlphaFold. While the TR regions of 17 of these proteins were unstructured, 26 were predicted to take up some secondary structure. In order to assess whether period length and copy number had any impact on the type of secondary structure attained by different TRs, a scatter chart was plotted (Figure 4). While TRs with period lengths 1-20 were seen to form structures with beta-pleated sheets, repeats with periods ranging from 20-40 formed either alpha helices or beta-pleated sheets, and the ones with periods longer than 40 contained alpha helices. Unstructured TRs were seen to be well distributed over all periods. Most structured TRs were seen to lie in the period range 20-40. While no relation could be established between copy number and secondary structures formed, TRs with short periods tended to have higher copy number.

**Figure 4:**
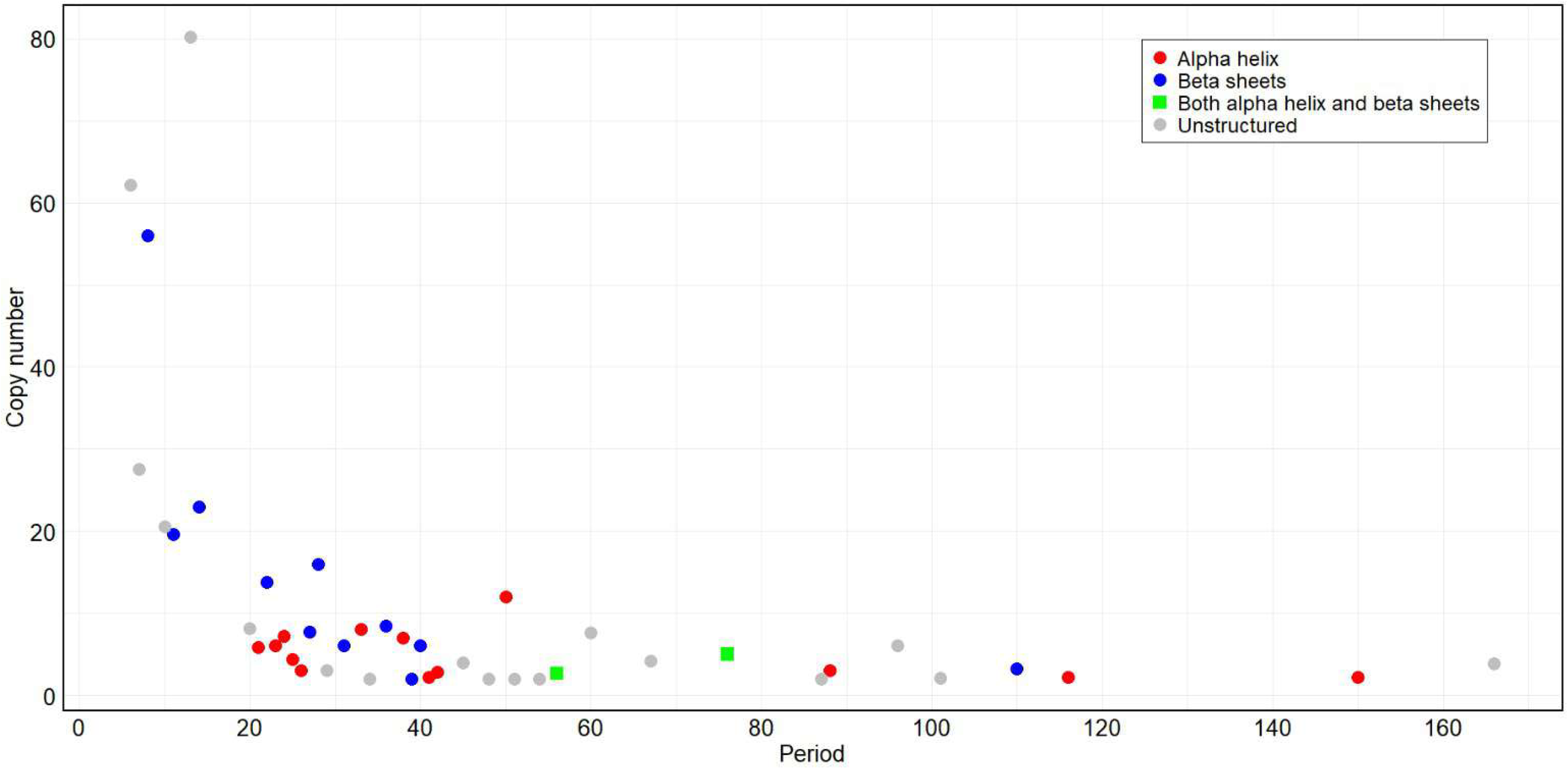
Secondary structure composition of repeat regions. Scatter plot showing relation of secondary structure with copy number and period of the repeat region. Each dot represents one repeat region with its period on the X axis and copy number on the Y axis. Secondary structure of repeat regions is denoted as follows: red dots: alpha helices, blue dots: beta pleated sheets, green squares: both alpha helices and beta pleated sheet, grey dots: unstructured.

**Table 5:** List of TR containing proteins for structural analysis. Table lists representative TR-containing proteins for different periods used for structure prediction and analysis. PlasmoDB ID, total protein length, start and end point of repeat region, period and copy number are tabulated.

| PlasmoDB ID | Annotation | Protein Length | Start | End | Period | Copy number |
| --- | --- | --- | --- | --- | --- | --- |
| PF3D7_1348400 | conserved Plasmodium membrane protein, unknown function | 3855 | 366 | 612 | **4** | 61.75 |
| PF3D7_0506500 | conserved Plasmodium protein, unknown function | 3980 | 3303 | 3509 | **5** | 41.4 |
| PF3D7_0424700 | serine/threonine protein kinase, FIKK family | 1222 | 559 | 933 | **6** | 62.17 |
| PF3D7_1318300 | conserved Plasmodium protein, unknown function | 1840 | 234 | 426 | **7** | 27.57 |
| PF3D7_1035200 | S-antigen | 585 | 105 | 552 | **8** | 56 |
| PF3D7_1038400 | gametocyte-specific protein | 9563 | 1306 | 1770 | **9** | 51.56 |
| PF3D7_0619000 | FPL domain-containing protein, putative | 1796 | 992 | 1197 | **10** | 20.6 |
| PF3D7_1035800 | probable protein, unknown function | 712 | 434 | 649 | **11** | 19.64 |
| PF3D7_0526600 | conserved Plasmodium protein, unknown function | 4524 | 1620 | 2028 | **12** | 34.08 |
| PF3D7_0201900 | erythrocyte membrane protein 3 | 2441 | 1324 | 2366 | **13** | 80.23 |
| PF3D7_1120000 | conserved protein, unknown function | 1070 | 115 | 437 | **14** | 23 |
| PF3D7_1446500 | nucleoporin NUP313, putative | 3001 | 1357 | 1536 | **15** | 12 |
| PF3D7_0403400 | zinc finger protein, putative | 3825 | 831 | 962 | **16** | 8.25 |
| PF3D7_1038400 | gametocyte-specific protein | 9563 | 923 | 1213 | **18** | 16.17 |
| PF3D7_1442600 | TRAP-like protein | 3543 | 1251 | 1560 | **19** | 16.32 |
| PF3D7_0208300 | conserved Plasmodium protein, unknown function | 1802 | 448 | 609 | **20** | 8.1 |
| PF3D7_1241900 | tetratricopeptide repeat protein, putative | 676 | 213 | 336 | **21** | 5.9 |
| PF3D7_0201900 | erythrocyte membrane protein 3 | 2441 | 753 | 1056 | **22** | 13.82 |
| PF3D7_1123500 | golgi protein 2 | 1336 | 269 | 407 | **23** | 6.04 |
| PF3D7_1308700 | conserved Plasmodium protein, unknown function | 607 | 416 | 589 | **24** | 7.25 |
| PF3D7_1201400 | Plasmodium exported protein, unknown function | 1129 | 836 | 944 | **25** | 4.36 |
| PF3D7_1128300 | ATP-dependent 6-phosphofructokinase | 1570 | 266 | 345 | **26** | 3.04 |
| PF3D7_1120600 | DDRGK domain-containing protein, putative | 2545 | 723 | 930 | **27** | 7.7 |
| PF3D7_0723300 | conserved protein, unknown function | 880 | 364 | 809 | **28** | 15.93 |
| PF3D7_1252800 | Plasmodium exported protein (PHISTb), unknown function | 626 | 410 | 497 | **29** | 3.03 |
| PF3D7_0526600 | conserved Plasmodium protein, unknown function | 4524 | 2971 | 3398 | **30** | 14.27 |
| PF3D7_1024800 | exported protein 3 | 1464 | 1132 | 1318 | **31** | 6.03 |
| PF3D7_0526600 | conserved Plasmodium protein, unknown function | 4524 | 811 | 972 | **32** | 4.75 |
| PF3D7_0500800 | mature parasite-infected erythrocyte surface antigen | 1434 | 573 | 837 | **33** | 8.03 |
| PF3D7_0701900 | Plasmodium exported protein, unknown function | 964 | 141 | 205 | **34** | 2 |
| PF3D7_1002100 | EMP1-trafficking protein | 631 | 237 | 541 | **36** | 8.47 |
| PF3D7_1038400 | gametocyte-specific protein | 9563 | 3323 | 3414 | **37** | 2.49 |
| PF3D7_1301200 | gametocyte-specific protein | 377 | 110 | 375 | **38** | 7 |
| PF3D7_0420600 | apicomplexan kinetochore protein 6, putative | 593 | 284 | 361 | **39** | 2 |
| PF3D7_1149100 | Plasmodium exported protein, unknown function | 450 | 127 | 367 | **40** | 6.03 |
| PF3D7_1001400 | exported lipase 1 | 921 | 256 | 346 | **41** | 2.22 |
| PF3D7_1201400 | Plasmodium exported protein, unknown function | 1129 | 629 | 750 | **42** | 2.88 |
| PF3D7_0526600 | conserved Plasmodium protein, unknown function | 4524 | 978 | 1092 | **43** | 2.67 |
| PF3D7_0526600 | conserved Plasmodium protein, unknown function | 4524 | 2782 | 2871 | **44** | 2 |
| PF3D7_0214300 | conserved Plasmodium protein, unknown function | 2500 | 484 | 663 | **45** | 4 |
| PF3D7_0508900 | protein AAP6 | 3134 | 2491 | 2638 | **47** | 3.02 |
| PF3D7_1024100 | ookinete maturation gene OMG1, putative | 1104 | 485 | 581 | **48** | 2.02 |
| PF3D7_1016300 | GBP130 protein | 824 | 220 | 821 | **50** | 12.04 |
| PF3D7_0214300 | conserved Plasmodium protein, unknown function | 2500 | 963 | 1064 | **51** | 2 |
| PF3D7_1038400 | gametocyte-specific protein | 9563 | 3138 | 3284 | **53** | 2.77 |
| PF3D7_1014300 | SPRY domain-containing protein, putative | 2379 | 950 | 1050 | **54** | 2 |
| PF3D7_1328200 | BRCA2 protein, putative | 2668 | 1340 | 1488 | **56** | 2.66 |
| PF3D7_0114000 | exported protein family 1 | 1463 | 316 | 773 | **60** | 7.6 |
| PF3D7_1442600 | TRAP-like protein | 3543 | 3130 | 3257 | **62** | 2 |
| PF3D7_0406500 | NYN domain-containing protein, putative | 3211 | 2315 | 2497 | **63** | 2.89 |
| PF3D7_1337500 | conserved Plasmodium protein, unknown function | 3334 | 2085 | 2203 | **64** | 2 |
| PF3D7_1237900 | conserved Plasmodium protein, unknown function | 1188 | 457 | 735 | **67** | 4.16 |
| PF3D7_1211800 | polyubiquitin | 381 | 1 | 380 | **76** | 5 |
| PF3D7_1035300 | glutamate-rich protein GLURP | 1233 | 499 | 672 | **87** | 2 |
| PF3D7_1035900 | merozoites-associated armadillo repeats protein | 566 | 180 | 443 | **88** | 3 |
| PF3D7_0730900 | EMP1-trafficking protein | 2110 | 1113 | 1692 | **96** | 6.04 |
| PF3D7_1038400 | gametocyte-specific protein | 9563 | 2673 | 2988 | **98** | 3.18 |
| PF3D7_0501400 | interspersed repeat antigen | 1720 | 391 | 603 | **101** | 2.11 |
| PF3D7_1038400 | gametocyte-specific protein | 9563 | 2161 | 2591 | **108** | 3.99 |
| PF3D7_0730900 | EMP1-trafficking protein | 2110 | 638 | 992 | **110** | 3.23 |
| PF3D7_1478600 | EMP1-trafficking protein | 1263 | 369 | 621 | **116** | 2.18 |
| PF3D7_1321100 | protein kinase domain-containing protein, putative | 3447 | 1124 | 1366 | **123** | 2 |
| PF3D7_1038400 | gametocyte-specific protein | 9563 | 6104 | 6486 | **124** | 3.08 |
| PF3D7_1321100 | protein kinase domain-containing protein, putative | 3447 | 839 | 1252 | **145** | 2.86 |
| PF3D7_0504700 | centrosomal protein CEP120, putative | 2349 | 1051 | 1384 | **150** | 2.23 |
| PF3D7_0501400 | interspersed repeat antigen | 1720 | 507 | 1140 | **166** | 3.82 |
| PF3D7_1038400 | gametocyte-specific protein | 9563 | 3493 | 3993 | **244** | 2 |
| PF3D7_1038400 | gametocyte-specific protein | 9563 | 6395 | 9017 | **250** | 10.46 |

**Table 6:** VAST result of selected TR-containing proteins. Table shows VAST results (top 3 hits) for predicted structures of GBP-130, MAAP, CERLI2 and EMP3. PlasmoDB ID (protein name), total length of repeat region, PDB ID_chain for the VAST hits, length of the structurally aligned region, VAST score, RMSD values and percent sequence identity for each TR containing protein are tabulated.

| Protein | PDB<br>_chain | Alignment<br>length | TR<br>length | VAST<br>Score | RMSD<br>(Å) | Sequence<br>identity (%) |
| --- | --- | --- | --- | --- | --- | --- |
| **GBP-130** | 4R10_A | 368 | 602 | 33.33 | 4.12 | 7.9 |
|  | 6FVB_A | 367 | 602 | 24.3 | 5.07 | 4.6 |
|  | 6KBM_A | 365 | 602 | 41.4 | 3.03 | 11 |
| **MAAP** | 6NOD_C | 191 | 264 | 19.73 | 1.32 | 12.6 |
|  | 3BWT_A | 191 | 264 | 22.25 | 2.06 | 11 |
|  | 3K49_E | 191 | 264 | 24.03 | 0.99 | 10.5 |
| **CERLI2** | 3VSV_D | 131 | 132 | 24.41 | 2.95 | 15.3 |
|  | 1QCX_A | 129 | 132 | 21.51 | 3.1 | 9.3 |
|  | 4C2L_A | 129 | 132 | 27.2 | 3.05 | 13.2 |
| **EMP3** | 6NYR_A | 301 | 304 | 22.72 | 0.39 | 24.6 |
|  | 7B1B_B | 300 | 304 | 21.51 | 3.2 | 9.3 |
|  | 4LXR_A | 299 | 304 | 21.6 | 3.18 | 8.7 |

Nearly two third of the repeat elements adopted alpha helical folds, and the rest were comprised mainly of beta pleated sheets. Several of TR regions were predicted by AlphaFold to form distinct super secondary structures with repetitive patterns (Figure 5). Many of these were open alpha or beta solenoids. None of the repeat regions were seen to attain a closed conformation. We used Swiss-Model to predict the structures of TR regions whose AlphaFold structures had very low to low confidence scores. PDB and UniProt IDs of the templates used for molecular modelling along with their percentage identity are tabulated in Table S4a. Our results showed erythrocyte membrane protein 3 (PF3D7_0201900), glycophorin binding protein GBP130 (PF3D7_1016300) and merozoites-associated armadillo repeats protein (PF3D7_1035900) to take up solenoid structures upon molecular modelling as opposed to long alpha helices depicted by AlphaFold. While PF3D7_1016300 and PF3D7_1035900 still had an alpha helical secondary structure, PF3D7_0201900 was now predicted to form beta helical solenoids.

**Figure 5:**
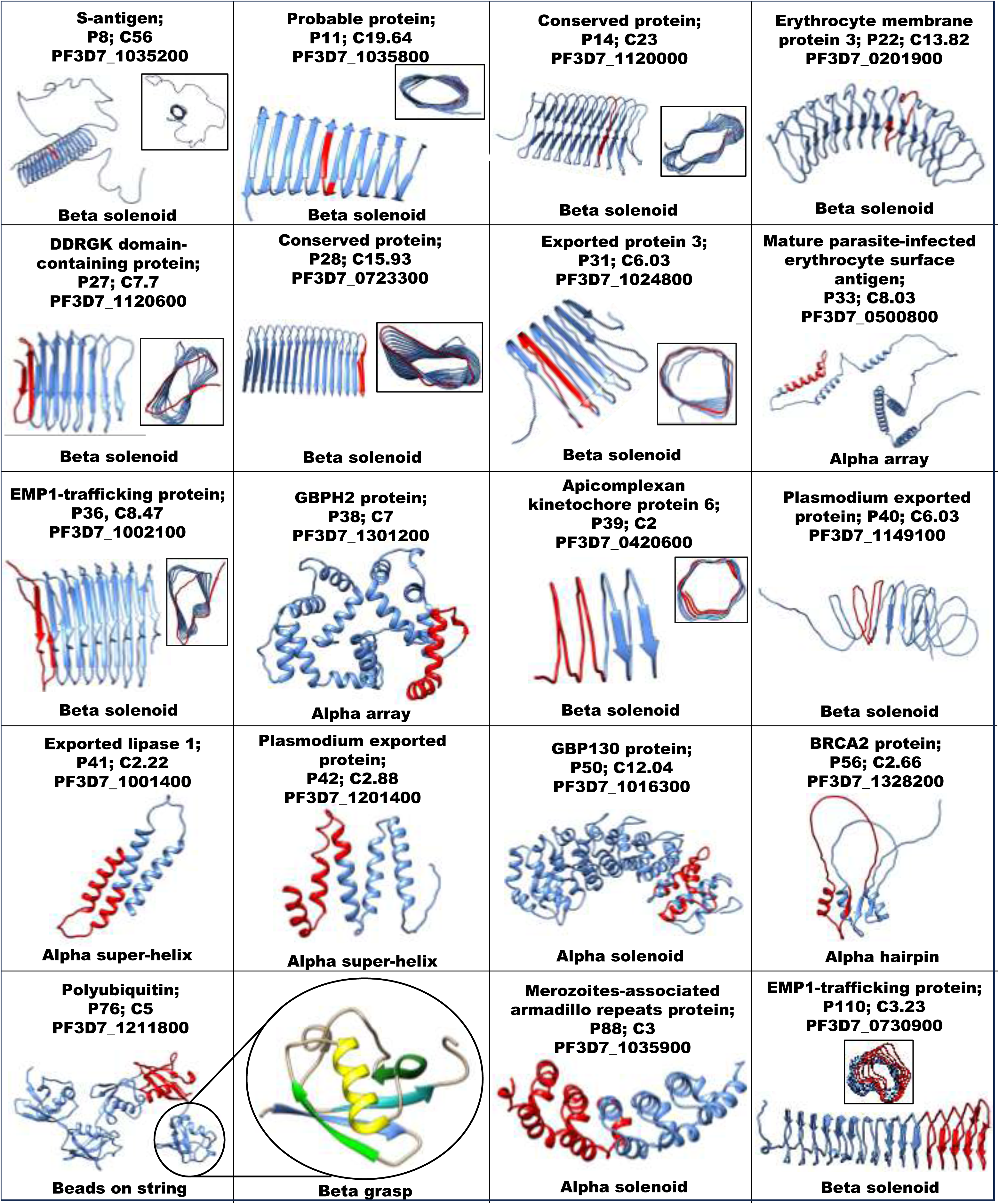
Representative super-secondary structures formed by repeat regions. Panel shows super-secondary structures formed by TR regions of 19 representative proteins. Names, PlasmoDB IDs, copy number and period are indicated above each ribbon diagram. Repeat unit is indicated in red colour. Cross-sectional views of a subset of structures are shown in the respective figures.

Several of the structured TR regions were seen to fold into beta helices, which are solenoid domains formed by a two/ three faced helical arrangement of parallel beta pleated sheets [87]. These included the repeat regions of S antigen (PF3D7_1035200), proteins of unknown function (PF3D7_1035800, PF3D7_0723300 & PF3D7_1120000), erythrocyte membrane protein 3 (PF3D7_0201900), DDRGK domain containing protein (PF3D7_1120600), exported protein 3 (PF3D7_1024800) and EMP1 trafficking proteins (PF3D7_1002100 & PF3D7_0730900). Beta helical structures often participate in cell-cell adhesion, interaction with partner proteins and other macromolecules like polysaccharides [87]. Other super secondary structures adopted by TR regions included various arrangements of alpha helices like alpha solenoids (PF3D7_1016300, PF3D7_1035900), alpha bundles (PF3D7_0500800), array (PF3D7_1301200, PF3D7_1201400) and alpha super helix (PF3D7_1001400). The merozoites-associated armadillo repeats protein (PF3D7_1035900) contained the armadillo (ARM) repeats which behave as ‘molecular Velcro’ having roles in protein-protein binding, cellular signalling, cell-cell adhesion, and regulation of cytoskeleton and transcription [88], [89]. There were only two super secondary structures that contained both alpha helices and beta pleated sheets. These were a putative BRCA2 protein (PF3D7_1328200) forming alpha hairpins and polyubiquitin (PF3D7_1211800). Polyubiquitin had a beads-on-a-string appearance where each bead was made of a beta grasp super secondary structure. This is a classical ubiquitin fold where 4 beta pleated sheets and an alpha helix arrange themselves to form a scaffold [90]. Interestingly, each of the above folds have been earlier reported to act as modules for intermolecular interactions, highlighting the ability of repetitive regions in proteins to carry out important biological functions.

### Repeat regions of proteins may be used to assess structure-function relationships

Structures taken up by a protein are ordinarily driven by their primary sequence. However, there are ample examples where different primary protein sequences adopt converging structural folds to perform similar functions [91]. We have therefore selected a few TR regions with striking repetitive patterns in their predicted super-secondary structures, and compared them with the PDB database by using NCBI Vector Alignment Search Tool (VAST) to identify structurally similar proteins [61]. VAST is a computer-based algorithm that follows geometrical principles to identify proteins with three-dimensional structures similar to the query, enabling recognition of structurally similar proteins without significant sequence identity. Structures and functions of the structurally similar proteins identified for each repeat region (top 3 hits with the highest VAST scores) were then assessed in an attempt to understand probable functions of these TR proteins.

The first protein chosen for understanding the structure-function relationship of repeat regions of Pf was Glycophorin-binding protein 130 (GBP-130; PF3D7_1016300; 824 aa long), which is a merozoite-surface protein that binds glycophorin receptor on host RBCs for invasion [92]. While the function of this protein has been reported, its structure has not been experimentally solved. Therefore, we used the predicted structure of the TR region of GBP-130 to understand its probable function and assess the correctness of our method. GBP-130 has a 50 mer consensus repeat sequence ‘ADNKEDLTSADPEGQIMREYAADPEYRKHLEIFHKILTNTDPNDEVERRN’ from aa residues 220 to 821 with a copy number of 12.04. Antibodies raised against a part of recombinantly expressed TR domain of PfGBP-130 were able to inhibit host cell invasion. While a minimum of 3.5-4.5 of these GBP-130 repeats were found to be necessary for glycophorin binding, it was reported that an increase in the number of repeats enhanced the binding [93]. Since the AlphaFold structure of the repeat region of GBP-130 showed very low (pLDDT <50) to low confidence (pLDDT <70), the structure of its TR region was modelled by Swiss-Model by using an engineered protein (PDB ID: 4RV1) as a template [94]. (Figure 6a, Table S4). pLDDT (predicted local distance difference test) values in AlphaFold are a per-residue measure of confidence in the built model that range from 0 to 100 i.e. Very high (pLDDT > 90), High (90 > pLDDT > 70), Low (70 > pLDDT > 50) and Very low (pLDDT < 50). NCBI VAST run using the modelled structure identified humpback-2 (HMP-2) protein of *C. elegans* (PDB ID: 4R10; Score: RMSD:) [95], Lph2, a novel bidirectional nuclear transport receptor in *S. cerevisiae* (PDB ID: 6FVB; Score: RMSD:) and vacuolar protein 8 (Vac8) (PDB ID: 6KBM; Score: RMSD:) [96] of *S. cerevisiae* as the three best hits based on their VAST score. All the above structures contained Armadillo (ARM) repeats, which have an arrangement of three helices per repeat unit [89], [97]. ARM repeats form an alpha solenoid when placed in tandem, and act as a platform for diverse cellular roles like cell adhesion and signalling through protein-protein interactions. Structural alignments of the TR domain of GBP-130 with the best VAST hits are shown in Figure S1. The central ARM domain of HMP-2, a homologue of mammalian beta-catenin, interact with cadherin HMR-1 to promote cell adhesion during body elongation in *C.elegans* [95]. Vac8 is a protein that performs multiple roles in nucleus-vacuole junction formation, autophagy etc. via its interaction with different proteins like Nvj1, Atg13 and Vac17 [96]. ARM repeat-containing proteins therefore perform very diverse functions with a central theme of protein binding that aligns well with the glycophorin binding activity of GBP-130.

**Figure 6:**
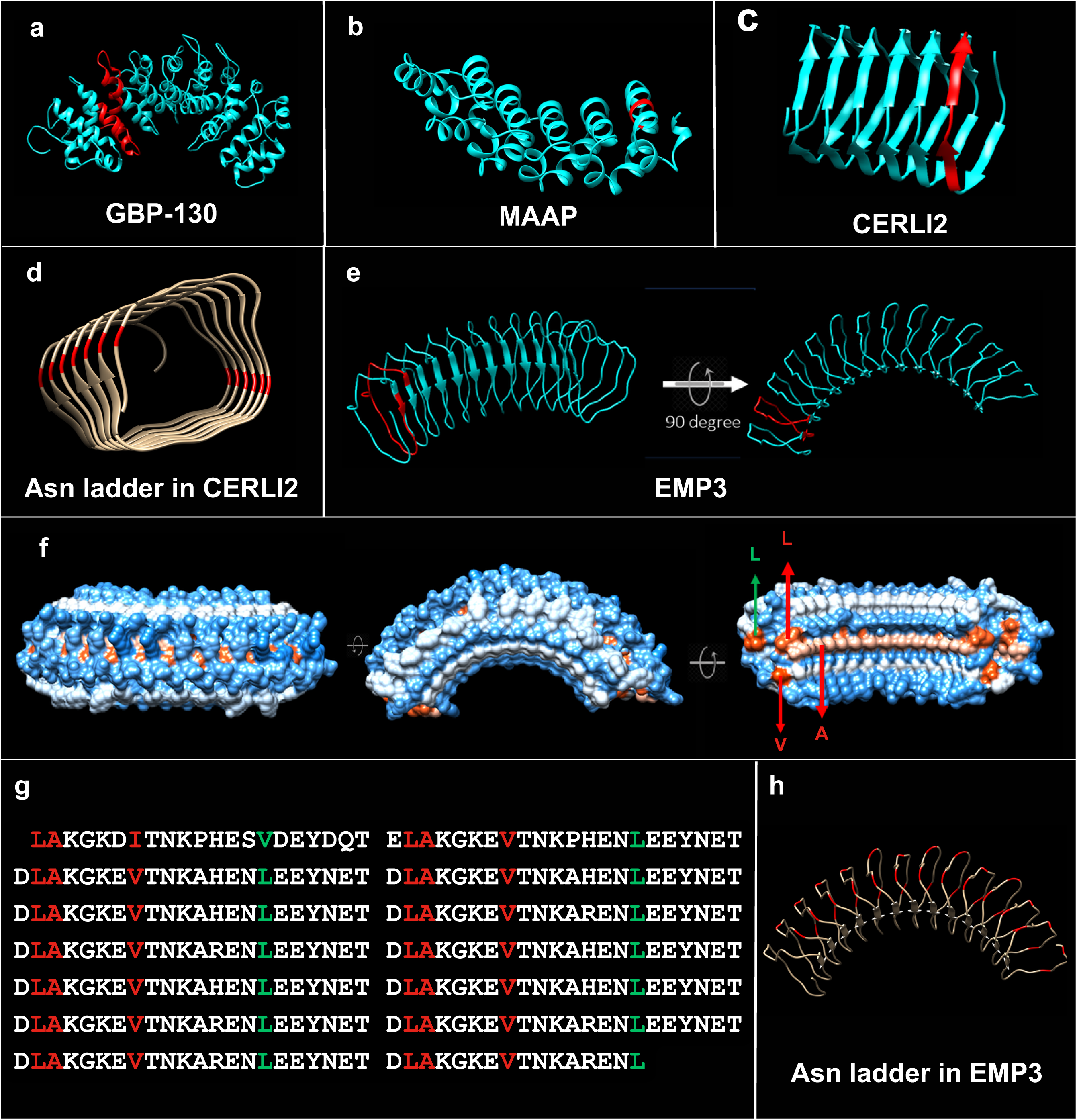
Analysis of predicted structures of selected repeat regions. Super secondary structures formed by a) GBP-130 b) MAAP c) CERLI2. Structures are shown as cyan ribbons, and one repeat unit is shown in red. d) Cross-sectional view of CERLI2 with Asn ladders (2 in number) highlighted in red. e) Ribbon diagram of EMP3 f) Hydrophobicity surface view of EMP3 showing hydrophobic residues in red and hydrophilic residues in blue. Amino acids L, L, V and A forming the hydrophobic backbone of the repeat units are labelled. g) Amino acid sequence of EMP3 highlighting residues forming hydrophobic backbone (as shown in figure 6g). h) Asn ladders of EMP3 (3 in number) highlighted in red.

Merozoite-associated armadillo repeats protein (MAAP; PlasmoDB ID: PF3D7_1035900; 566 aa long) is a Pf protein expressed in late erythrocytic stages of the parasite (merozoites and schizonts) with a role in host cell invasion [88]. A TR region of PfMAAP spanning residues 180 to 443 (period: 88; copy number: 3) contains ARM repeats that form an open alpha-solenoid. Since the AlphaFold structure showed a very low to low confidence score, the structure of the above TR domain of PfMAAP was also modelled by using the structure of Puf domain of Pumilio homolog 2 from mice (PDB ID: 3GVO; percent identity: 11.52%) (Figure 6b, Table S4). NCBI-VAST identified *C. elegans* PUF-8 (PUmilio/FBF-8) complexed with RNA (PDB ID: 6NOD), an RNA-binding domain of PUF-1 from *S. cerevisiae* (PDB ID: 3BWT) and RNA bound PUF-3 (PDB ID: 3K49) from *S. cerevisiae* to have structural similarity with the modelled structure (structural alignments shown in Figure S1). PUF-8 is an RNA binding protein involved in organism development, spermatogenesis and sexual reproduction of *C. elegans* [98]. PUF-1 and PUF-3 are also RNA binding proteins involved in regulation of cellular processes, translation, macromolecular metabolic processes and post-transcriptional regulation [99], [100]. Domain analysis by using InterPro predicted PfMAAP to carry a pumilio-homology domain (PUM-HD) from 202 to 556 that overlaps with its repeat region [101], [102]. PUM-homologs are RNA binding proteins that are crucial for germ cell maintenance, regulation of stem cell fate, development and neurological functions [103], [104], [105], [106]. GO analysis of PfMAAP also predicts this protein to have RNA binding properties [52].

Pf cytoplasmic exported rhoptry leaflet interacting protein 2 (PfCERLI2) (PlasmoDB ID: PF3D7_0405200) is a rhoptry bulb protein that has an essential role in merozoite invasion, maintenance of rhoptry morphology, and antigen processing [107]. Rhoptries are a pair of club-shaped organelles located at the apical tip of merozoites [108]. These play a critical role in binding and penetration of host erythrocytes by merozoites through secretion of important proteins during the invasion process. PfCERLI2 is believed to have arisen from an ancestral gene duplication of cerli1 that encodes PfCERLI1 (PF3D7_0210600; sequence identity 24.65% with query cover 31%). Both these proteins carry the conserved ‘PHIS motif’ with consensus sequence ‘PHIS[-]xxP’ and 2 conserved domains i.e. Pleckstrin homology (PH) and C2. In addition to the functions performed by PfCERLI2, PfCERLI1 is also known to have roles in antigen secretion from rhoptries. PfCERLI 2, a 579 amino acid long protein is longer than PfCERLI1 (446 amino acids) due to addition of a repeat containing domain at its C terminal end (435 to 566). This TR region has the 10 mer consensus sequence ‘EIKNDHIQTD’ repeating 13.2 times. The predicted structure of the TR region of PfCERLI2 on AlphaFold was used to identify similar structures by using NCBI VAST, and found to form a right-handed parallel beta helix. Figure 6c shows a ribbon diagram of the TR domain of PfCERLI 2. This super secondary structure has four beta pleated sheets per turn of the beta helix, which is a characteristic feature of the polygalacturonase family of enzymes [109]. Right-handed parallel beta-helices have an internal core made up of hydrophobic amino acids and homostacks of polar, aliphatic or aromatic amino acids within their core to maintain structural stability [110], [111]. The structure of PfCERLI2 TR is stabilized by the presence of Asparagine (Asn) ladders that form hydrogen bonds between the side chain amides (Figure 6d). Asn ladders are a common feature of parallel beta helices and repetitive modules like leucine-rich repeats, amyloid fibrils and toll-like receptors [87]. The three structures identified by NCBI VAST to be most similar to the TR region of PfCERLI 2 were those of Xylan-1,4-beta-xylosidase (XylC) (PDB ID: 3VSV Chain D; RMSD: 2.95), pectin lyase B (PDB ID: 1QCX; RMSD: 3.10) and a xylogalacturonan hydrolase (PDB ID: 4C2L; RMSD: 3.05) [112], [113], [114]. The structure of XylC was seen to have three domains i.e. two right-handed parallel beta-helices (residues 1-75 and 201-638) and a beta-sandwich (residues 76-200). Xyl C is a hydrolase that breaks glycosidic bonds in xylose. Figure S1 shows the alignment of the TR region of PfCERLI2 with the second right-handed parallel beta-helix of XylC that is bound to xylose via hydrogen bonding at its active site, an open carbohydrate binding cleft. Pectin lyase B and xylogalacturonan hydrolase are also hydrolytic enzymes that cleave alpha-1,4-glycosidic bonds of galacturonate moieties (structural alignments shown in figure S1). It is therefore likely that the repeat region of PfCERLI 2 is involved in carbohydrate binding and hydrolysis in the malaria parasite *P. falciparum*.

Another protein that showed a striking repetitive pattern was *P. falciparum* erythrocyte membrane protein 3 (PfEMP3; PlasmoDB ID: PF3D7_0201900). PfEMP3 is a 300 kDa protein (2441 amino acids) that resides intracellularly associated with the membrane of infected red blood cells [115]. Targeted mutants of PfEMP3 have been earlier shown to have defective cytoadherence, knob formation and PfEMP1 trafficking. These mutants were formed from deletion of parts of a C-terminal repeat region of PfEMP3 (13 mer consensus sequence ‘QQNTGLKNTPSEG’ repeating 80.23 times from amino acids 1324 to 2366). Also, the N-terminal region of PfEMP3 (38-97) has been reported to bind with the erythrocyte cytoskeletal protein ‘spectrin’ [116]. Here, we have analysed the structure-function relationship of a different repeat unit in the central region (753 - 1056) of PfEMP3, which has the 22 mer consensus sequence ‘LAKGKEVTNKAHENLEEYNETD’ repeating 13.82 times. While Alphafold predicts this TR region to be an alpha helix, the confidence levels of the predicted structure were very low (pLDDT <50). Also, its Ramachandran plot analysis and structure validation by ERRAT was poor (Figure S2a). Therefore, we performed molecular modelling of this region by using SwissModel where CroV588 from the virus *Cafeteria roenbergensis* (PDB ID; 23.26% sequence identity; query cover) was used as a template (Table S4). Here, the repeat region was predicted to form a curved beta solenoid (one-faced parallel beta helix) made up of beta pleated sheets and coils (Figure 6e). Secondary structure prediction using JPred also showed that the repeat region of PfEMP3 is likely to form beta pleated sheets and coils, and not alpha helices (Figure S2b) [117]. The 13 beta pleated sheets (formed by 3 repetitive residues D/E-L-A) in the predicted structure are parallel to each other and linked through coils that are 19 amino acids long. The curved solenoid has a hydrophobic backbone formed by two conserved leucine residues, an alanine and a valine (Figure 6f-g). The coils in this structure are stabilized by three distinct Asn ladders (homostacks of Asparagine) (Figure 6h). The best hit identified by NCBI VAST to be most structurally similar to the TR region of PfEMP3 was CroV588 (PDB ID: 6NYR Chain A; RMSD:) [118], the viral protein of unknown function that had also been used as a template for molecular modelling. Subsequent hits with a high score were the cytokine receptor Toll 5A of Aedes aegypti (PDB ID: 7B1B Chain B; RMSD:) [119], toll receptor ectodomain from *Drosophila melanogaster* (PDB ID: 4LXR Chain A; RMSD:) [120] and toll-like receptor 3 (TLR3) from *Mus musculus* (PDB ID: 7C77 Chain A; RMSD:) [121] (Table S4). While toll receptors in insects have roles in developmental patterning and innate immunity through their binding with Spatzle cytokines, toll-like receptors of higher organisms act as pattern recognition receptors (PRRs) to elicit immune responses via the toll signaling pathways. Toll receptors and TLRs sense DNA and RNA for controlling innate immunity [122]. A characteristic feature of all the above structures is that these are receptors that form an arc shaped beta solenoid, a common component of virulence factors, toxins, allergens etc [87]. Figure S1 shows the structural alignment of the TR region of PfEMP3 with its topmost VAST hits.

## Conclusion

Tandem repeats are a common feature of many eukaryotic proteins that hold a wealth of information and are poorly characterized. TRs often contribute to organism biology as either a structural component or functional entity of a protein. Pf proteins have been earlier reported to be rich in homopolymeric tracts of Asn, which is a feature associated with protein aggregation. Our study provides a comprehensive evaluation of *Plasmodium* proteomes to identify their oligomeric and polymeric repeat content termed ‘repetome’, and a detailed analysis of tandem repeats in *P. falciparum*. Our work highlights the potential of repetome mining as a generalized strategy to uncover the hidden information in the repeat regions of varied proteomes, using Pf as an example. TRs in a protein are often linked with the ability to interact with other macromolecules efficiently due to the increased avidity provided by multiple copies of a binding motif [17]. Repetitive regions may therefore act as interaction hubs by binding with other proteins/ molecules allowing functional talk and complexation [123]. The exceptional richness of TRs in Pf and Pv may be evolutionarily driven for improved virulence *via* host-pathogen interactions, implementation of immune evasion strategies, dampening of the host immune response *etc*. Structure elucidation or prediction for the TR domains of Pf in a proteome-wide manner alongside building structure-function relationships in the future is likely to provide glimpses into the missing links between *Plasmodium* biology and disease pathogenesis.

## Abbreviations

TR: Tandem Repeats
Pf: *Plasmodium falciparum*
PEXEL: *Plasmodium* export element
PfEMP1: *Plasmodium falciparum* erythrocyte membrane protein 1
STRP: Structured tandem repeat protein
EC number: Enzyme Commission Number
LCR: Low complexity region
VSA: Variant surface antigen
iRBC: Infected red blood cell
PDB: Protein Data Bank;
ARM: Armadillo repeat motif

---

**Figure S1:**
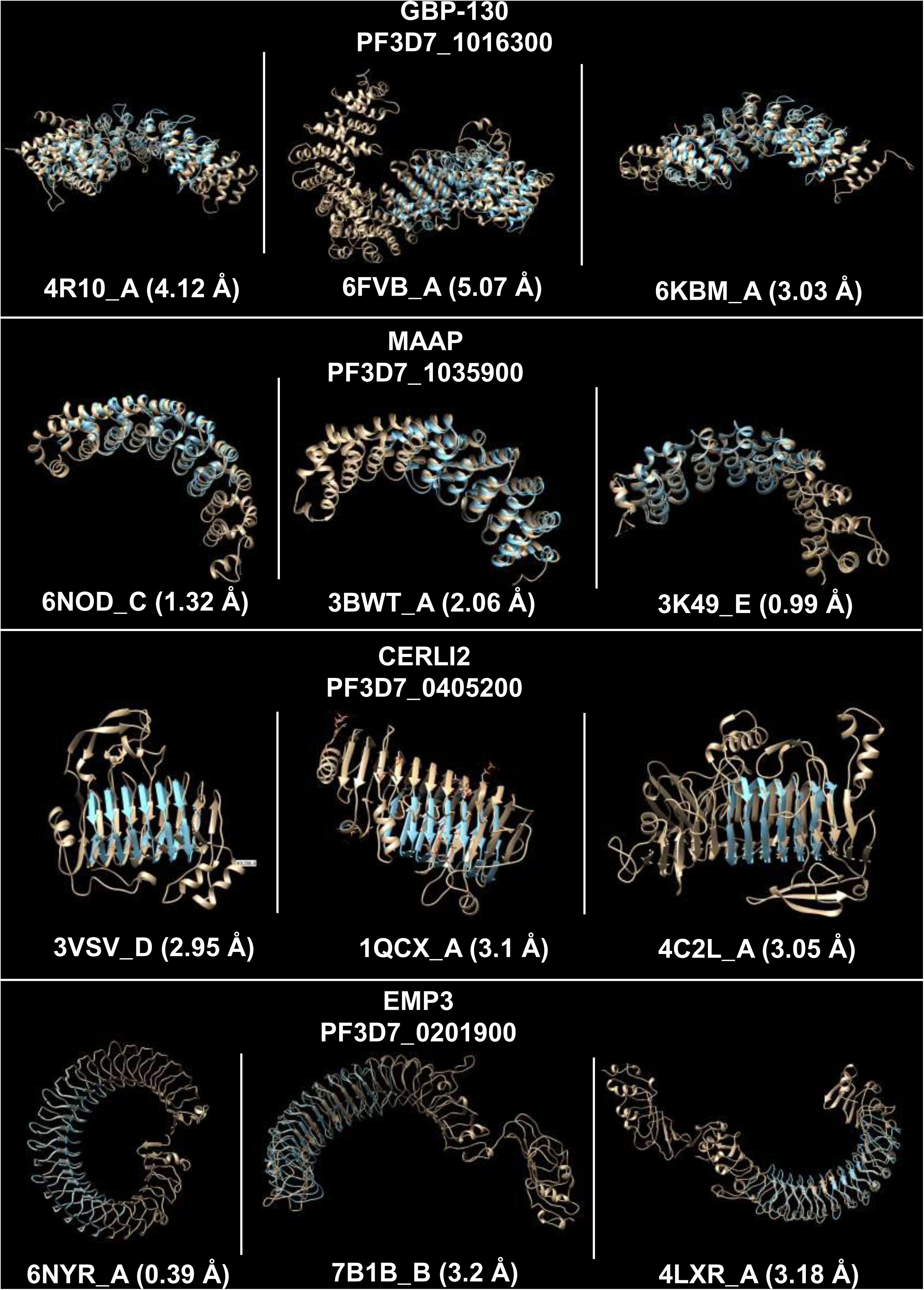
Structure alignment of repeat regions with VAST hits. Figure shows alignment of predicted structures of GBP-130, MAAP, CERLI2 and EMP3 with experimentally determined structures of their top three VAST hits. The structure of repeat regions is shown in cyan and those of VAST hits in beige colour. PDB IDs_chain for each VAST hit are given below each aligned structure.

**Figure S2:**
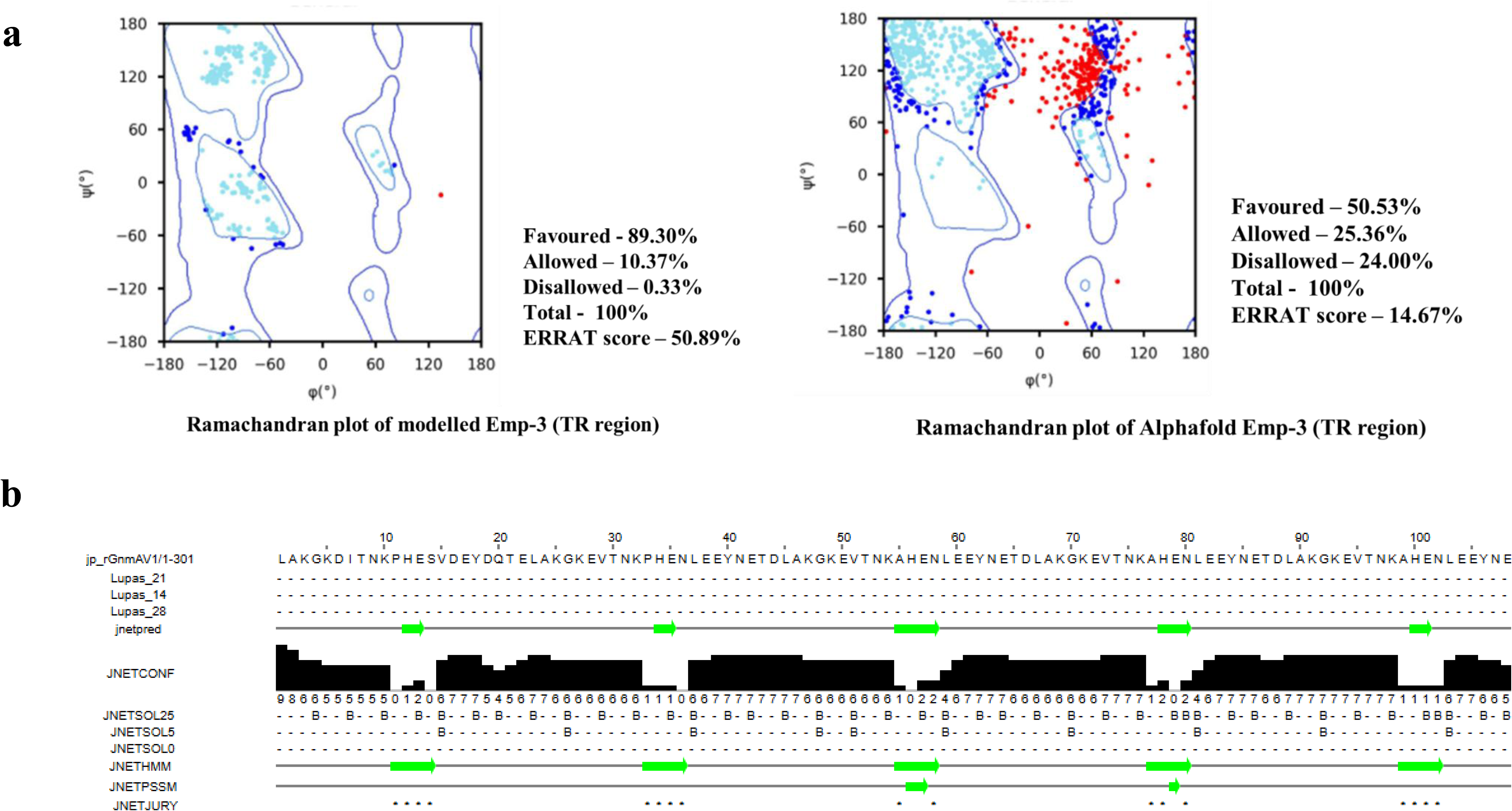
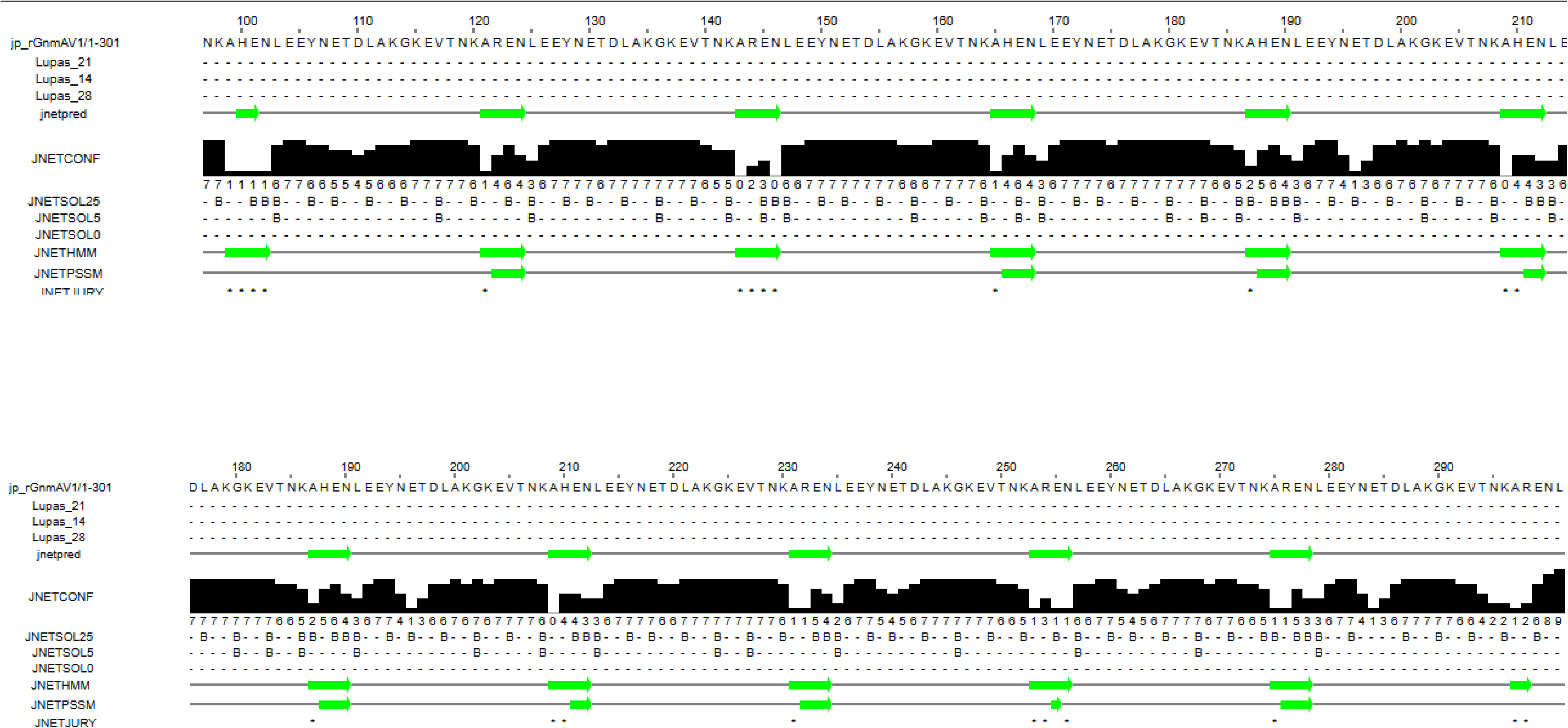
Ramachandran plot and ERRAT analysis of EMP3: a) Figure compares Ramachandran plot analysis and ERRAT for Alphafold and predicted structure (SwissModel) b) Secondary structure prediction by JPred

**Table S1:** Cellular localization and annotation of TR containing Pf proteins. Table lists TR containing Pf proteins having >50% of their length in the repeat region. PlasmoDB IDs, annotation, sequence coverage in TR region expressed as percentage, presence/ absence of PEXEL motif and their cellular localization reported on plasmodb.org is tabulated.

| PlasmoDB ID | Annotation | TR percentage | PEXEL | Location |
| --- | --- | --- | --- | --- |
| PF3D7_1211800 | Polyubiquitin | 99.74 | Absent | cell surface;cytoplasm;nucleus;ubiquitin ligase complex |
| PF3D7_1038400 | Gametocyte-specific protein (Pf11-1) | 89.26 | Absent | integral component of membrane; host cell cytoplasm |
| PF3D7_0501400 | Interspersed repeat antigen (FIRA) | 84.43 | Absent | Maurer's cleft;extracellular space |
| PF3D7_1035200 | S-antigen | 76.58 | Absent | symbiont-containing vacuole;vacuole |
| PF3D7_0201900 | Erythrocyte membrane protein 3 (EMP3) | 74.59 | Present | integral component of membrane; Maurer's cleft;cytoplasm;extracellular vesicle;keratohyalin granule;nucleus;symbiont-containing vacuole |
| PF3D7_0909500 | Subpellicular microtubule protein 1, putative (SPM1) | 73.23 | Absent | cell surface;cytoskeleton;subpellicular microtubule |
| PF3D7_0730900 | EMP1-trafficking protein (PTP4) | 73.08 | Present | integral component of membrane; host cell;nucleus |
| PF3D7_1016300 | GBP130 protein (GBP130) | 73.06 | Present | integral component of membrane; host cell cytoplasm;infected host cell surface kAbsentb;nucleus |
| PF3D7_0831800 | Histidine-rich protein II (HRP2) | 72.80 | Absent | Maurer's cleft;cytoplasm;food vacuole;host cell cytoplasm;host cell membrane |
| PF3D7_1301200 | GBPH2 protein (GBPH2) | 70.56 | Present | integral component of membrane; host cell |
| PF3D7_1149100 | Plasmodium exported protein, unkAbsentwn function | 66.29 | Present | host cell |
| PF3D7_1036400 | Liver stage antigen 1 (LSA1) | 63.86 | Absent | integral component of membrane; actin filament bundle assembly |
| PF3D7_1401000 | GBPH protein (GBPH) | 63.84 | Present | host cell |
| PF3D7_1372200 | Histidine-rich protein III (HRPIII) | 62.17 | Present | Maurer's cleft;food vacuole |
| PF3D7_0702300 | Sporozoite threonine and asparagine-rich protein (STARP) | 55.92 | Present | cell surface |
| **PF3D7_0723300** | Conserved protein, unkAbsentwn function | 55.69 | Absent | inner membrane pellicle complex;nucleus |
| **PF3D7_1102300** | Plasmodium exported protein, unkAbsentwn function | 53.24 | Absent | integral component of membrane; host cell periphery |
| **PF3D7_0108500** | ELM2 domain-containing protein, putative | 53.07 | Absent | nucleus; Maurer's cleft;nucleus |
| **PF3D7_1035900** | Merozoites-associated armadillo repeats protein (MAAP) | 52.83 | Absent | apical part of cell;cell surface; entry into host |
| **PF3D7_1035800** | Probable protein, unkAbsentwn function (M712) | 50.85 | Absent |  |
| **PF3D7_0114000** | Exported protein family 1 (EPF1) | 50.17 | Absent | integral component of membrane; Maurer's cleft |

**Table S2:**
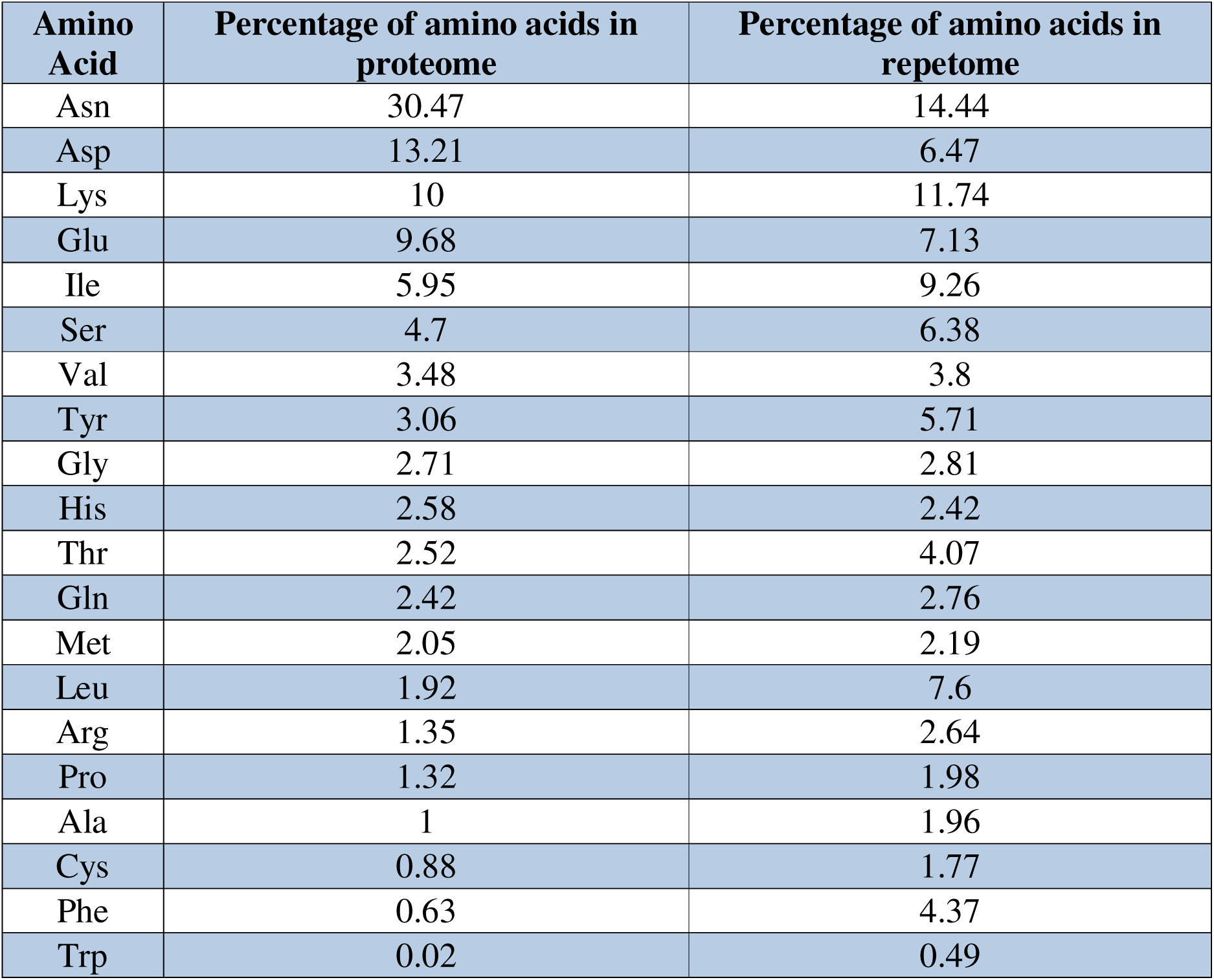
Amino acid composition of Pf Repetome vs Pf proteome. The percentage of 20 standard amino acids in Pf repetome and Pf proteome are tabulated.

**Table S3:**
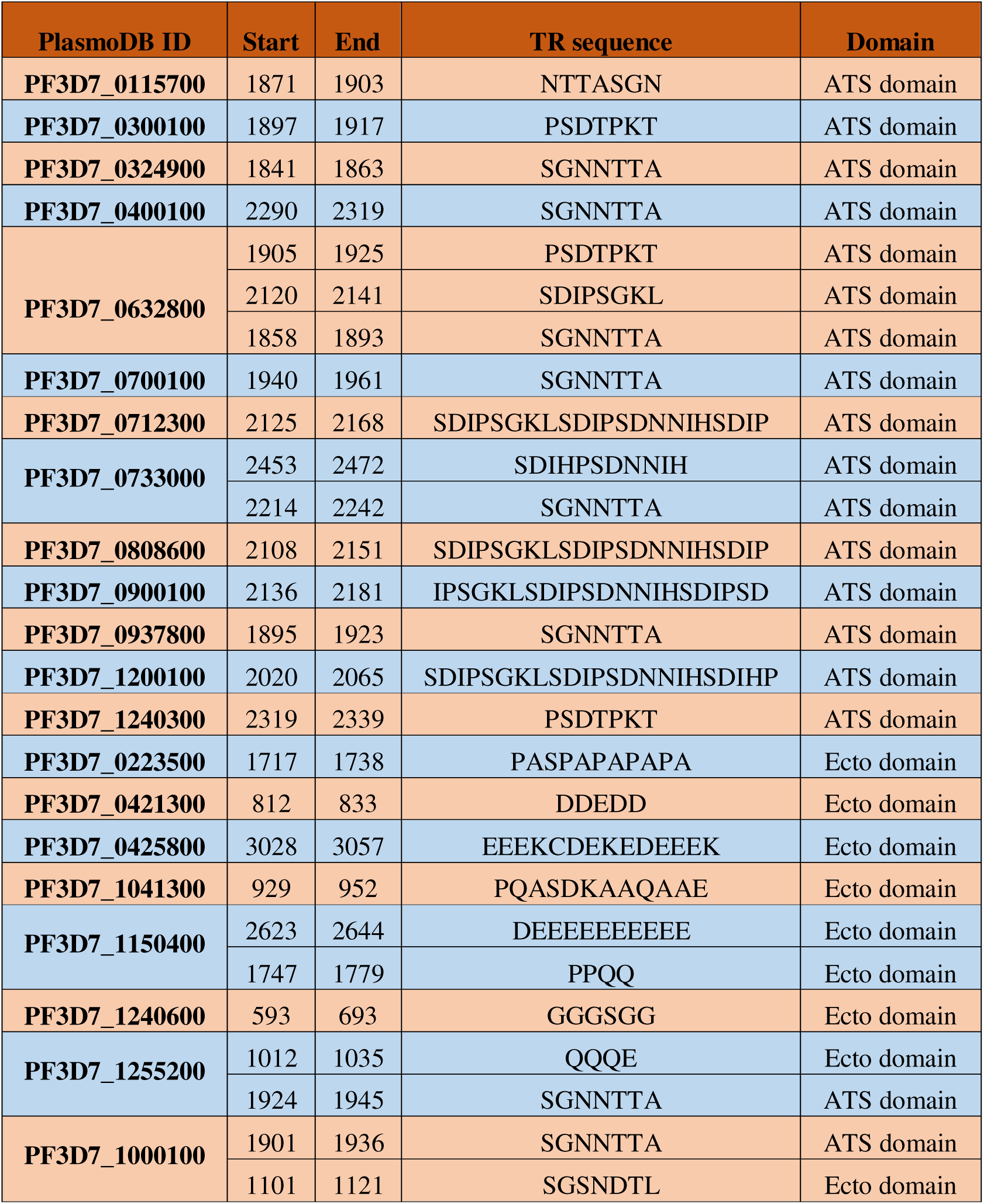
Repeats of PfEMP1 proteins. Table lists 21 TR containing PfEMP1 proteins. Consensus sequence of each repeat, repeat region boundaries and domain containing the repeat are tabulated.

**Table S4:** Molecular modelling. Table lists the Pf TR containing proteins whose structures were predicted by SwissModel. PlasmoDB IDs (protein name), PDB ID of template, percentage sequence identity of TR region with template, percentage of amino acids in allowed region (Ramachandran plot analysis) and ERRAT scores of the modelled structure are tabulated.

| Protein | EMP3 | GBP-130 | MAAP |
| --- | --- | --- | --- |
| Template PDB ID_chain | 6NYR_A | 4RV1_A | 3GVO |
| Sequence identity (%) | 23.26% | 33.69% | 11.52% |
| Ramachandran plot analysis |  |  |  |
| Favoured | 89.30% | 88.01% | 97.88% |
| Allowed | 10.37% | 9.96% | 1.06% |
| Disallowed | 0.33% | 2.04% | 1.06% |
| Total | 100% | 100% | 100% |
| ERRAT Overall Quality Factor | 50.89% | 85.97% | 93.44% |

